# Cognate Antigen Engagement Induces HIV-1 Expression In CD4^+^ T Cells From People On Long-Term ART

**DOI:** 10.1101/2024.02.27.582096

**Authors:** Milica Moskovljevic, Filippo Dragoni, Nathan L. Board, Fengting Wu, Jun Lai, Hao Zhang, James R. White, Rebecca Hoh, Kenneth Lynn, Pablo Tebas, Karam Mounzer, Steven G. Deeks, Luis J. Montaner, Janet D. Siliciano, Francesco R. Simonetti, Robert F. Siliciano

**Author notes:** Correspondence to: Francesco R. Simonetti, M.D., Ph.D., Virus Immunity and Pathogenesis Center, Department of Medicine, Johns Hopkins University School of Medicine, 855 N Wolfe Street, Baltimore, MD, 21205, USA., Robert F. Siliciano, M.D., Ph.D., Department of Medicine, Johns Hopkins University School of Medicine, 733 N Broadway, Baltimore, MD, 21205, USA. Francesco R. Simonetti and Robert F. Siliciano contributed equally to this work.

## Abstract

Despite antiretroviral therapy (ART), HIV-1 persists in latently-infected CD4^+^ T cells, preventing cure. Antigens drive the proliferation of infected cells, precluding latent reservoir decay. However, the relationship between antigen recognition and HIV-1 gene expression is poorly understood since most studies of latency reversal use agents that induce non-specific global T cell activation. Here, we isolated rare CD4^+^ T cells responding to cytomegalovirus (CMV) or HIV-1 Gag antigens from participants on long-term ART and assessed T cell activation and HIV-1 RNA expression upon co-culture with autologous dendritic cells (DCs) presenting cognate antigens. Physiological presentation of cognate antigens induced broad T cell activation (median 42-fold increase in CD154^+^CD69^+^ cells) and significantly increased HIV-1 transcription (median 4-fold), mostly through the induction of rare cells with higher viral expression. Thus, despite low proviral inducibility, physiologic antigen recognition can promote HIV-1 expression, potentially contributing to spontaneous reservoir activity on ART and viral rebound upon ART interruption.

## Introduction

Antiretroviral therapy (ART) effectively blocks viral replication, but HIV-1 persists in a small pool of latently-infected resting CD4^+^ T cells that harbor inducible, replication-competent proviruses^1–3^. This latent reservoir is the major barrier to cure. The reservoir was originally characterized with a quantitative viral outgrowth assay (QVOA) in which resting CD4^+^ T cells, which are non-permissive for viral replication, are stimulated with agents that cause polyclonal T cell activation, including the mitogen phytohemagglutinin (PHA) and the combination of anti-CD3 and anti-CD28 antibodies^1,2,4^. T cell activation induces changes in the transcriptional environment that allow induction of latent proviruses. These changes include nuclear translocation of the transcription factors NF-κB and NFAT, which bind the HIV-1 LTR and promote initiation of transcription^5–9^. The QVOA has been used to demonstrate the extremely slow decay rate of the reservoir^10–12^, which requires persons living with HIV (PWH) to maintain lifelong adherence to ART. Interruption of ART leads to rapid viral rebound, presumably due to the activation of latently-infected cells^13,14^. Preventing rebound from the latent reservoir is a major goal of HIV-1 cure research, but the nature of the stimuli that induce latent HIV-1 in vivo to cause rebound remains unclear. One hypothesis is that encounter with cognate antigen (Ag) is the critical event leading to rebound. This is a difficult hypothesis to test because the Ag-specificity of latently infected cells is largely unknown.

Although the role of Ag stimulation in viral rebound is unclear, there is now strong evidence that Ag plays a key role in the persistence of the latent reservoir. The reservoir persists not through new infection events, but rather through the long-term survival and proliferation of infected cells^15–23^. Potential factors driving the proliferation of HIV-1-infected cells include encounter with cognate or cross-reacting antigens^20,24,25^, as well as homeostatic cytokines such as IL-7 and IL-15^26^, and effects related to rare HIV-1 integration events into genes involved in cell survival or proliferation^18,19,27^. Repeated exposure to Ags derived from persistent viral infections, such as HIV-1 and cytomegalovirus (CMV), can drive extensive clonal expansion of infected CD4^+^ T cells with T cell receptors specific for the relevant antigens^20,24,28^. This clonal expansion may be partly responsible for the recent observation that the frequency of latently infected cells appears to plateau or even increase in people who have been on ART for more than 20 years^12^.

While chronic Ag exposure can drive clonal expansion of infected cells in vivo, the role of Ag in latency reversal is less clear. In vitro studies with polyclonal T cell activators indicate that cellular activation can lead to proviral induction and productive infection. However, productively infected cells die rapidly due to viral cytopathic effects and immune clearence^29,30^ as well as activation-induced cell death (AIDC)^31^. Therefore, HIV-1-infected cells responding to recurrent Ag stimulation could be progressively lost. However, the majority of infected cells in PWH on long-term ART are found in expanded clones^20^, suggesting some degree of uncoupling between T cell activation and HIV-1 expression. Previous work has shown that some proviruses cannot be induced even after multiple rounds of maximal mitogen stimulation ex vivo^32^, but the relationships between Ag-mediated T cell activation, HIV-1 gene expression, and cell proliferation remain unclear.

The study of HIV-1 gene expression in infected, Ag-responding cells is particularly challenging. First, the frequency of infected CD4^+^ T cells in peripheral blood and lymph nodes is low, typically less than 1 in 1000 cells, and the cells harboring inducible and intact HIV-1 are even less frequent^33,34^. Second, the frequencies of memory T cells responding to any particular Ag are also low, mostly ranging between 0.001 to 5%^35,36^. As a result, most studies of HIV-1 latency reversal rely on ex vivo activation of CD4^+^ T cells by stimuli that induce polyclonal T cell activation^37,38^. These stimuli may not reproduce physiologic CD4^+^ T cell activation driven by Ag presented on professional antigen presenting cells such as dendritic cells (DC). Ag-driven activation involves a complex interaction between an Ag-derived peptide presented by a major histocompatibility complex (MHC) molecule (denoted as pMHC) and the T cell receptor (TCR). The TCR may behave as a mechanosensor^39–41^, explaining how the cell can be triggered by recognition of a very small number pMHC complexes on a DC. Polyclonal stimuli acting through the TCR bypass this mechanosensing mechanism and may induce different transcriptional responses. Previous studies of Ag-driven latency reversal^42,43^ have not used isolated Ag-responding cells and therefore cannot distinguish between direct Ag-mediated latency reversal and bystander T-cell activation. Here, to investigate the impact of Ag recognition on induction of HIV-1 gene expression in latently infected CD4^+^ T cells, we employed the ex vivo enrichment and expansion of CMV and HIV-1 Gag-reactive CD4^+^ T cells from people on ART. This approach allowed us to directly test whether Ag presented in a physiologic manner on autologous dendritic cells (DCs) could reverse HIV-1 latency.

## Results

### Enrichment and expansion of Ag-responding cells

To understand whether physiologic encounter with pMHC complexes on autologous DCs can cause latency reversal in Ag-responding CD4^+^ T cells, we isolated CD4^+^ T cells reactive to Ags derived from chronic viral pathogens, specifically CMV or HIV-1 itself. We enrolled 10 participants who had been on suppressive ART for a median of 11 years with no evidence of ongoing cycles of viral replication. Eight participants had undetectable HIV-1 RNA in plasma, while participants P6 and P9 had persistent and intermittent non-suppressible viremia (see Table S1 for participant characteristics). To isolate Ag-responding cells, between 4x10^7^ and 9x10^7^ CD8-depleted peripheral blood mononuclear cells (PBMCs) were stimulated with either lysates of CMV-infected cells, with HIV-1 Gag protein, or peptides pools for 18 hours, as previously described^20,44^. CMV- or Gag-responding CD4^+^ T cells were then isolated using two sequential column enrichment steps with magnetic beads conjugated with anti-CD154 (CD40L) antibodies^45^ (Figure 1A). CD154 is transiently expressed on the surface of CD4^+^ T cells following encounter with Ag. This method has been used previously in the characterization of CD4^+^ T cell responses to commensal and pathogenic fungi and SARS-CoV-2^36,44,46,47^.

**Figure 1.**
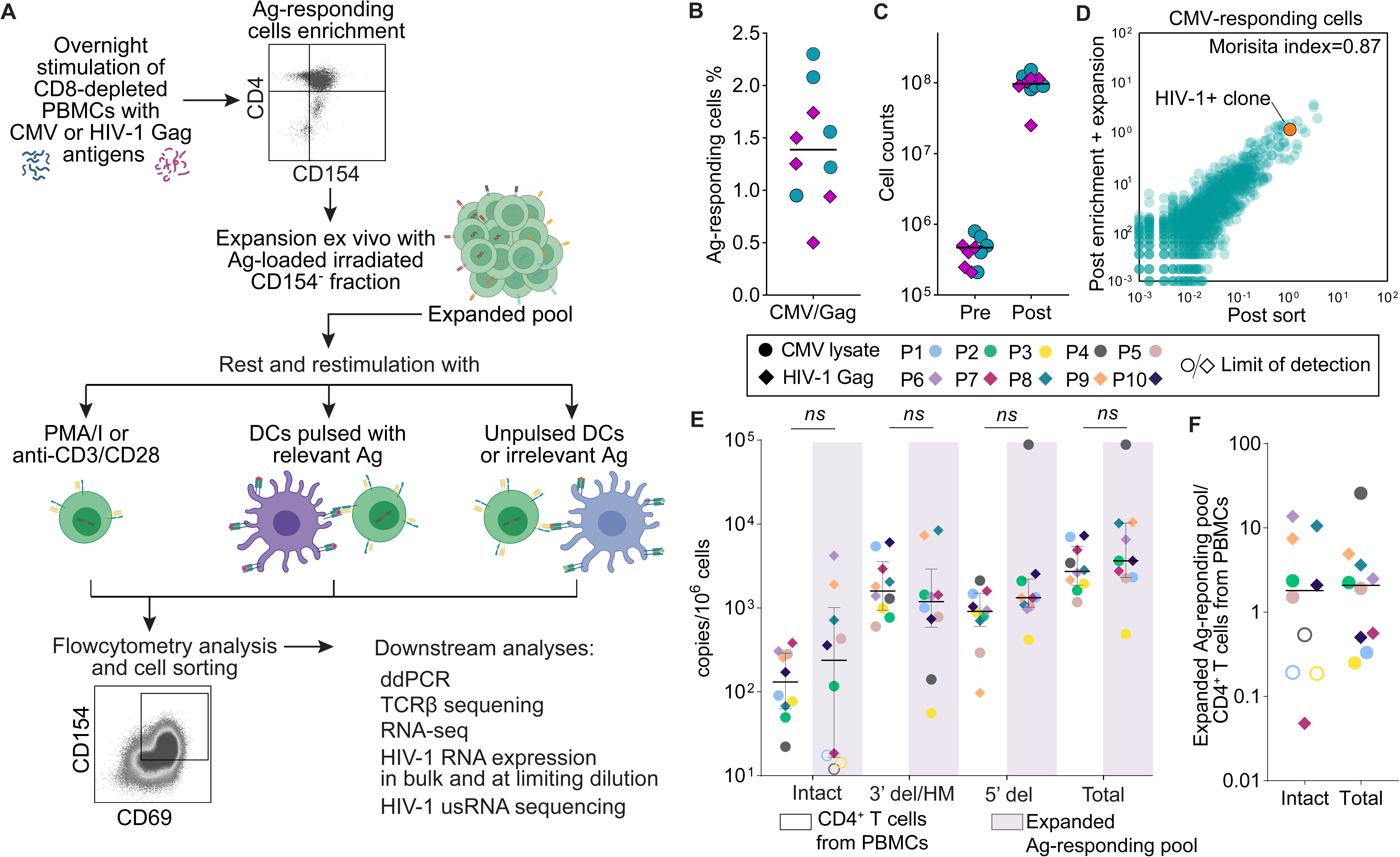
Experimental approach to study Ag-mediated T cell activation and induction of HIV-1 gene expression. (**A**) Schematic of experimental design. Enrichment of Ag-responding cells based on magnetic beads coated with anti-CD154 antibodies, and subsequent expansion ex vivo. Expanded cells were rested and subjected to either non-specific T cell activators (anti-CD3/CD28 activation beads or PMA/I) or cocultured with autologous monocyte-derived dendritic cells (DCs) loaded with either cognate Ag (CMV lysates or HIV-1 Gag peptides), neoantigen (KLH), or no addition of antigen (NoAg). (**B**) Percentage of Ag-responding cells among CD8-depleted PBMC estimated based on CD154 magnetic enrichment. (**C**) Cell count of Ag-responding cells before and after expansion. (**D**) Distribution of TCRβ sequences among Ag-responding cells from participant P1 at the end of the enrichment and expansion protocol compared to sequences obtained by overnight Ag stimulation of CD8-depleted PBMC and direct sorting of CD69^+^CD154^+^ T cells. Abundance of TCRβ sequences of Ag responding cells after Ag-specific expansion compared to direct sorting after overnight Ag-stimulation; representative scatter plot from participant P1. Circles indicate percentage abundance of individual productive TCRβ sequences in the two samples; the orange dot represents a well characterized clonotype carrying a defective provirus integrated into the *MKL1* gene^20^. Data on sorted cells are from a sample obtained 3 years prior to the current study^20^ (Simonetti et al, 2021) (**E**) Quantification of intact proviruses, proviruses with 3′ deletions and/or hypermutation (3′del/HM), proviruses with 5′ deletions (5′del), and total proviruses in purified, unstimulated CD4^+^ T cells and in the expanded pool of Ag-responsive cells. Total proviruses were calculated as the sum of intact, 3′del/HM, and 5′del proviruses. Horizontal bars represent median with interquartile range. Significance was determined by paired t-test. (**F**) Intact and total proviruses within expanded Ag-responding pools expressed as the fold change relative to purified CD4+ T cells isolated from PBMCs. Each dot represents a value from one study participant. Open symbols indicate values below the limit of detection. Horizontal bars represent median values.

Cells from participants P1-P5 were stimulated with CMV antigens because of their known co-infection with CMV and the previous characterization of their CMV-responding cells (P1-P4^20^), while cells from participants P6-P10 were stimulated with HIV-1 Gag as they had readily detectable CD4^+^ T cell responses to Gag in preliminary screening experiments. CMV- or Gag-responding cells were detected in samples from all 10 selected participants. The frequencies of Ag-responding cells, estimated based on cell input and column yields, were 0.5-2.3% (Figure 1B). This is consistent with values previously determined for the same donors by flow cytometric analysis of activation-induced marker (AIM) expression following stimulation with the relevant Ags^20^. Enriched cell populations were then expanded ex vivo. The CD154^-^ cells from the magnetic enrichment were irradiated and used as antigen-loaded cells to bolster the proliferation of Ag-responding cells for 10-14 days, as previously described^44^ (see Methods). Antiretroviral drugs (10 nM dolutegravir (DTG), 10 μM emtricitabine (FTC), and 10 μM tenofovir (TDF)) were included in the culture media to prevent de novo infection events. The expansion of Ag-responding cells resulted in a >2 log_10_ increase in cell number (Figure 1C), allowing us to characterize the cells and their response to subsequent stimulation. Since cell expansion ex vivo could skew the distribution of clonal populations, we sequenced the T cell receptor β chain (TCRβ) repertoire of the expanded cell pools, and compared TCRβ frequencies to those of Ag-responding cells directly sorted from PBMCs of the same participants based on upregulation of CD154 and CD69 in a previous study^20^. As shown in Figure 1D, the hierarchy of the most expanded clones of Ag-responding cells and the overall clonal population patterns were preserved, indicating that the TCRβ repertoires were stable over time in vivo and not significantly distorted by the expansion in culture (Figures 1D and S1A). In participant P1, the Morisita index^48^ – a metric of sample overlap– was high between AIM^+^ sorted cells and those obtained by our enrichment plus expansion protocol. The extent of TCRβ overlap was also comparable between the cells sampled immediately after CD154^+^ enrichment, after expansion, and by direct sorting from previous time points (Figure S1A). Similarly, analysis of TCRβ sequences for P2 and P3 showed strong correlation of clonal frequencies between AIM+ sorted cells and enriched cells expanded ex vivo (Figure S1B). Together, these results demonstrate that the enrichment and expansion protocol does not noticeably change the repertoire of Ag-responding cells.

### Frequency and composition of proviruses within expanded Ag-responding cells

To measure the frequency of HIV-1 infected cells among total and expanded Ag-responding CD4^+^ T cells, we used the intact proviral DNA assay (IPDA) ^49^. The IPDA is a droplet digital PCR assay that allows separate quantification of defective and potentially intact proviruses^49^. Prior to enrichment and expansion, the frequency of cells carrying intact proviruses varied from 22 to 384 per 10^6^ total CD4^+^ T cells (median 131, Figure 1E), consistent with previous studies^49,50^. As expected, the frequencies of intact proviruses were substantially lower than those of total proviruses (1174-7271 copies per 10^6^ CD4^+^ T cells, median 2723), reflecting the small fraction of proviruses that are intact (median 3.5%, Figures 1E and S1C). Among Ag-responding cells expanded ex vivo, the overall frequencies of cells with either intact or defective proviruses were not significantly different from the frequencies in CD4^+^ T cells from PBMCs (Figure 1E). Intact proviruses were also highly variable across pools of expanded cells (median 238, range 12-4205 copies/10^6^ cells). For example, in P6 and P9 the frequency of intact proviruses was >10^3^ copies/10^6^ cells, likely due to dominant clones carrying intact proviruses among Gag-responding cells. We did not detect intact proviruses within expanded CMV-responding cells in 3 out of 5 participants (Figure 1E). This may suggest that intact proviruses in some participants (P1, P3 and P4) were either rare among CMV-responding cells or did not proliferate as well as uninfected cells during the expansion, consistent with recent observations that cells carrying intact proviruses may proliferate less upon stimulation ex vivo^51^. Although the differences in the proviral frequencies between expanded Ag-responding cells and total CD4^+^ T cells from PBMCs were highly variable, the median ratio values were close to 2 (1.8 and 2.1 for intact and total proviruses, respectively, Figure 1F). Taken together, these results demonstrate the striking heterogeneity of proviral composition within Ag-responding cells.

### Re-stimulation with antigen-loaded dendritic cells induces T cell activation

At the end of expansion ex vivo, cells were washed and rested in basal media for three days in the presence of antiretroviral drugs. After rest, we observed downregulation of the transient activation markers CD25 and CD69, but some cells continued to express HLA-DR (Figure S2A). These results suggest that the cells returned to a partially quiescent state, allowing us to characterize their response to cognate Ag and polyclonal activation.

In vivo, CD4^+^ T cell responses to Ag are often initiated by contact with dendritic cells (DCs) presenting processed forms of the relevant Ag^52^. To test the Ag-reactivity in a physiologic manner, we generated autologous, monocyte-derived DCs loaded with Ags of interest (CMV-lysate or HIV-1 Gag peptides and Gag^p55^ protein) or with the irrelevant Ag Keyhole limpet hemocyanin (KLH). Rested populations of enriched, Ag-responding CD4^+^ T cells were co-cultured with autologous, Ag-pulsed DCs. In addition, CD4^+^ T cells were cultured with DCs that had not been pulsed with any external Ag (NoAg). As positive controls, CD4^+^ T cells were treated in parallel with polyclonal activating stimuli including anti-CD3/CD28 beads (1:1 cell to bead ratio), PMA/Ionomycin (PMA/I), or the protein kinase C (PKC) agonist bryostatin, a well-studied LRA^53^. As a negative control, some rested cell were left untreated (NoTx). After 18 hours, cells from all conditions were analyzed by flow cytometry. We assessed activation of CD4^+^ T cells by surface expression of CD69 and CD154 on live cells (Figure 2A and Figure 2B). Although there was some basal expression of CD69 in untreated cells and a modest increase by co-culture with unpulsed DCs, CD69 was significantly upregulated by coculture with cognate Ag-loaded DCs (Figures 2A, 2B and S2B). Upregulation of CD69 by Ag-loaded DCs was similar to that induced by PMA/I and anti-CD3/CD28 (median fold increase in median fluorescence intensity (MFI) relative to NoTx was 6.78, 9.61, and 5.60, respectively, p=0.001) (Figure S2B). Expression of the activation-induced costimulatory molecule CD154 was significantly increased by polyclonal or Ag-specific stimulation (Figures 2A, 2B and S2C). Of note, although the upregulation of CD69 and CD154 induced by PMA/I and CD3/CD28 was variable, it was not significantly different from stimulation with cognate antigens (Figures 2C and 2D). Importantly, autologous DCs pulsed with the relevant Ag caused an even higher fraction of cells expressing CD154 and CD69 (median 61, range 50-79%) compared to PMA/I (median 53, range 6-82%) (Figure S2D). In addition, Gag peptide pools and whole recombinant Gag^p55^ recombinant proteins led to comparable levels of T cell activation (Figure 2D), suggesting that Gag-reactive cells recognized epitopes as a result of antigen processing and presentation. No increase in T cell activation was seen following exposure to DCs pulsed with KLH, confirming the specificity of the cells obtained at the end of enrichment and expansion. Stimulation with bryostatin did not cause significant increase in CD69^+^CD154^+^ CD4^+^ T cells (median 5.75%, Figure S2D). Overall, these results indicated that the enrichment and expansion method yield a high fraction of CD4^+^ T cells that can be activated by re-exposure to their cognate Ag.

**Figure 2.**
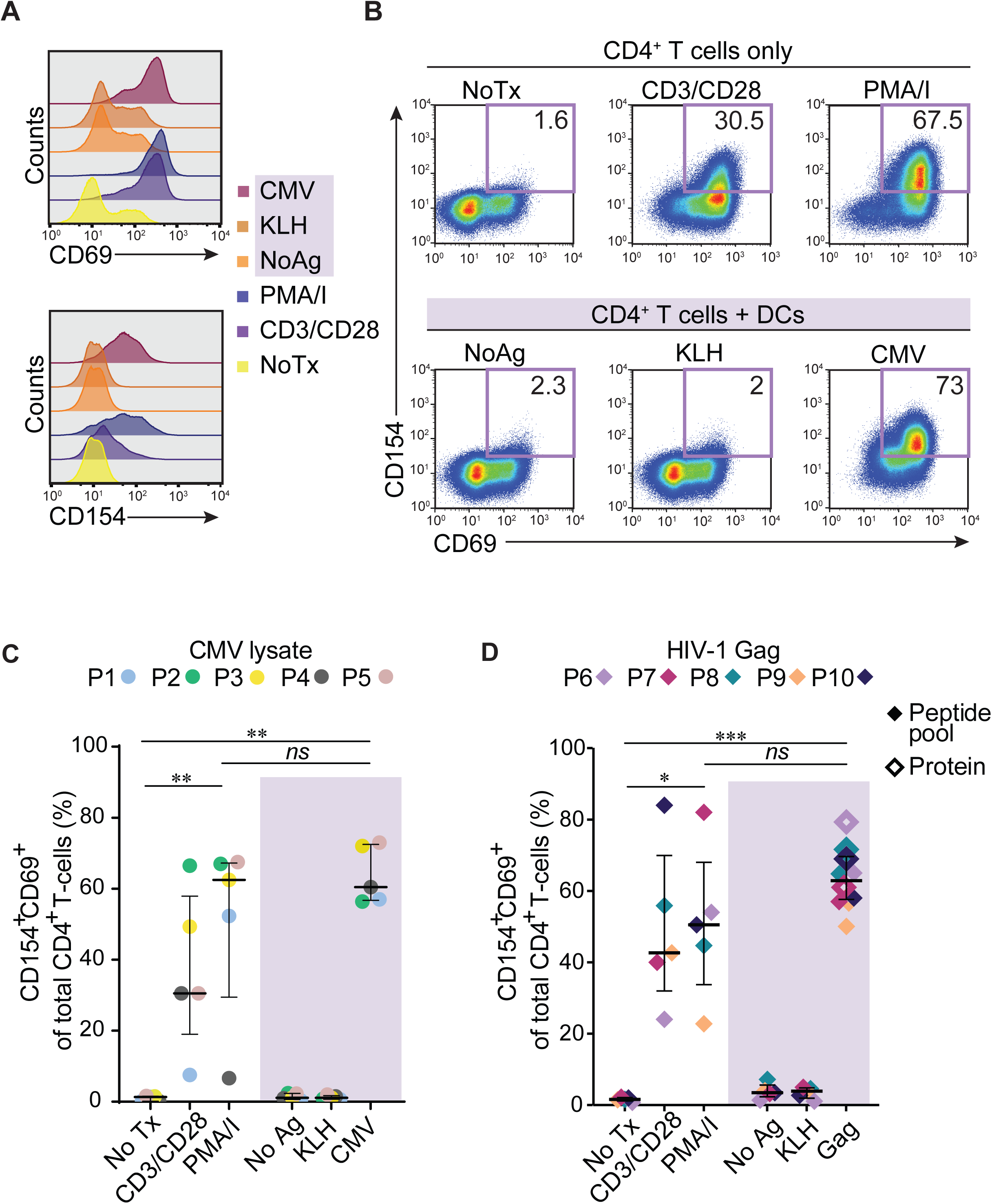
Activation of Ag-responding cells by exposure to different stimuli. (**A**) Representative histograms for CD69 and CD154 surface expression following 18 hours incubation of resting Ag-responding cells. Results are shown for one representative participant (P5). Shaded region of the figure legend represents coculture with DCs. (**B**) Representative flow cytometry plots showing upregulation of CD154 and CD69 on CD4^+^ T cells from a representative CMV-responding participant (P5). Top row: response to no treatment (NoTx) or polyclonal activating stimuli. Bottom row: response to DC pulsed with no Ag, an irrelevant Ag (KLH), or with CMV lysate. (**C)** Percentage of C154^+^CD69^+^CD4^+^ T cells in expanded pool of CMV-responding cells (left panel) and (**D**) Gag-responding cells (right panel) following an 18 hour incubation with the indicated stimuli. Horizontal bars represent median with interquartile range. Significance was determined by one-way ANOVA followed by Kruskal Wallis’s test for multiple comparisons, *P<0.05, **P<0.01, ***P<0.001.

### Encounter with cognate Ag engages pathways critical for HIV-1 reactivation

Although Ag-pulsed DCs and non-specific stimulation led to a comparable increase of CD69^+^CD154^+^ cells, we hypothesized that the stimuli that bypass the physiologic Ag recognition mechanism mediated by the α and β chains of the TCR or the early steps of TCR signal amplification (e.g. ZAP70) would induce distinct signatures in transcriptional activation, which in turn could impact HIV-1 gene expression^41^ (Figure 3A). To address this hypothesis, we performed bulk mRNA-sequencing on live CD4^+^ T cells from 6 participants after 18 hours of culture with the following conditions: no treatment (NoTx), unpulsed DCs (NoAg), cognate Ag-pulsed DCs, anti-CD3/CD28, and PMA/I (Figure 3A, see methods). Global transcriptome analysis showed clear clustering of samples based on culture conditions, with the NoTx and NoAg conditions giving expression patterns that were clearly distinct from those seen with activating stimuli (cognate Ag, CD3/CD28, and PMA/I). However, there were clear differences in patterns of gene expression induced by cognate antigen and those induced by polyclonal stimuli (Figure 3B). To specifically investigate downstream signaling, we narrowed the transcriptome analysis to genes whose promoters are targeted by the transcription factors NF-κB and/or NFAT. These two transcription factors regulate effector programs upon T cell activation and bind to the cis-acting elements of the HIV-1 LTR, driving viral gene expression^5,6,54–56^. We compiled a list of 199 genes (125 for NF-κB and 89 for NFAT) based on transcription factor-promoter binding data and review of the literature (see methods and Figure S3A). Principal component analysis (PCA) based on this gene set showed that CD4^+^ T cells clustered by the type of stimulation, even among those causing T cell activation (Figure 3C). Interestingly, stimulation with cognate antigens led to distinct clustering of CMV- and Gag-responding cells. Th17 and Th22 effector molecules were among the most variable genes (*IL17A*, *IL17F*, *IL22*), likely reflecting the different composition of polarized subsets between CD4 responses induced by CMV and HIV-1^25,57^. We next assessed differential gene expression between cells stimulated with DCs loaded with cognate Ags, and non-specific stimulation with anti-CD3/CD28 and PMA/I (Figure 3D and 3E and Figures S3B and 3SC). While there were only few differentially expressed genes between cells treated with cognate Ag and anti-CD3/CD28 (6 and 3 NF-κB-regulated genes (Figure 3D), and 2 and 7 NFAT-regulated genes (Figure S3B), respectively) we observed increased expression of multiple NF-κB and NFAT target genes with PMA/I stimulation (Figures 3E and S3C). These genes included effector molecules such as cytokines (*IL2*, *TNF*, *CFS2*[GM-CSF], *IFNG*, *IL17*) and chemokines (*CCL3*, *CCL4*), and genes involved in cell survival and division (*BCL2L1*, and *BCL2A1*, *CCND1*)^58,59^. Conversely, cognate Ags and anti-CD3/CD28 showed increased expression of genes involved in regulatory responses (*IL10*, *IL4*, *IDO1*, *FOXP3*) and immune exhaustion (*HAVCR2*, *TOX*).

**Figure 3.**
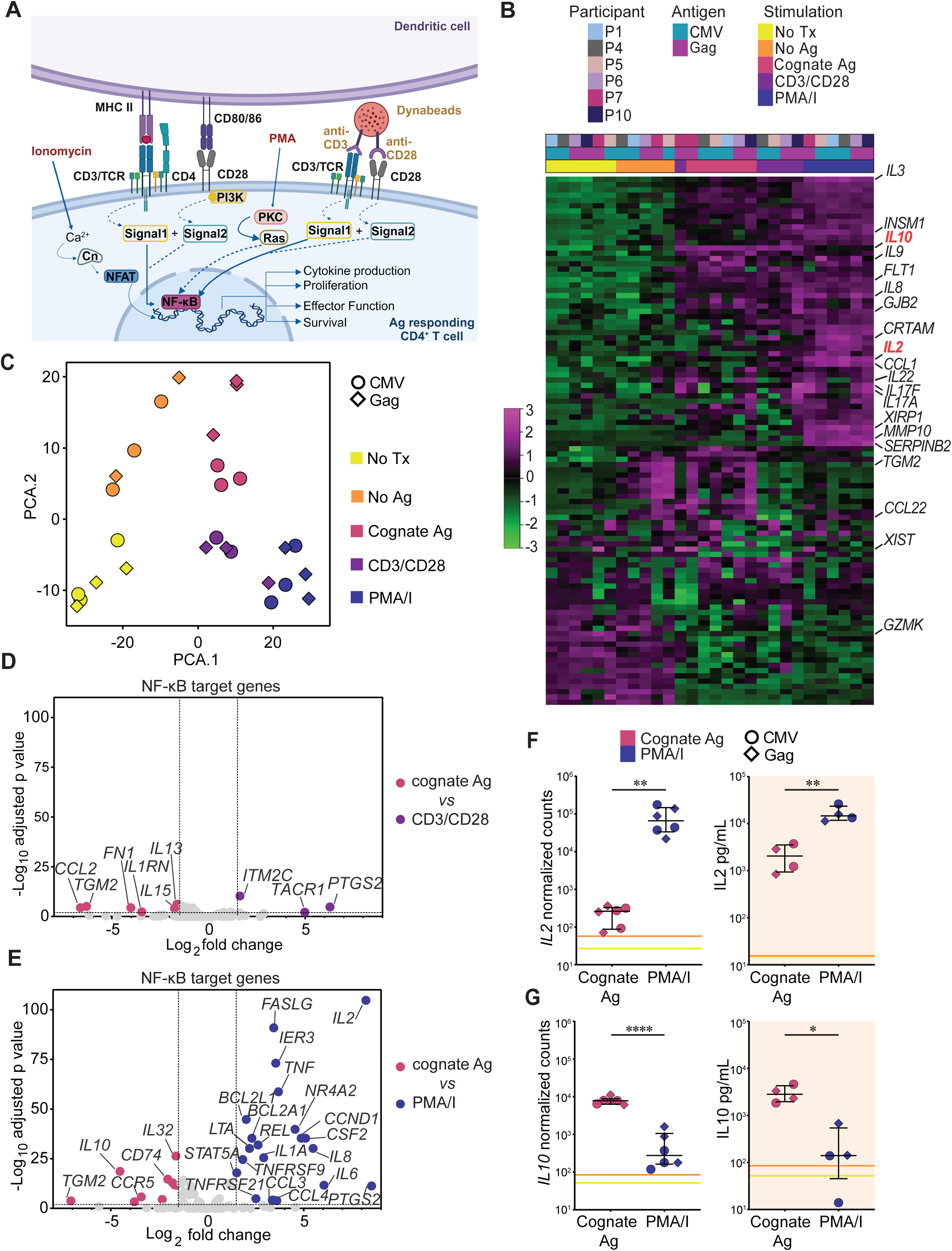
Distinct transcriptional activation in Ag-responding CD4^+^ T cells upon encounter with cognate Ag compared to PMA/I stimulation. (**A**) Schematic of TCR signaling network upon CD4^+^ T cell activation through Ag-presentation on the surface of DCs, anti-CD3/CD28 or PMA/I. (**B**) Heatmap of gene expression and hierarchical clustering based on the Pearson correlation of the 100 most variable genes; Z-score cut off was set at 4; the top 20 genes are labelled; scale bar indicates normalized relative expression. (**C**) Principal component analysis (PCA) based on gene expression of NF-κB and NFAT target genes. (**D**) Volcano plot of differentially expressed genes (DEGs) in cognate Ag versus anti-CD3/CD28 for NF-κB target genes (**E**) Volcano plot of DEGs NF-κB target genes in cells stimulated with cognate Ag versus PMA/I. (**F**) Normalized counts of *IL2* expression (left) and supernatant protein levels of IL2 (right) in cells stimulated with cognate Ag versus PMA/I.(**G**) Normalized counts of *IL10* expression (left) and supernatant levels of IL10 (right) in cells stimulated with cognate Ag versus PMA/I. (**F-G**) Each symbol corresponds to a study participant. Horizontal bars represent median with interquartile range. All data shown as average of technical replicates. Statistical significance was determined by unpaired t-test*P<0.05, **P<0.01, ***P<0.001, ****P<0.0001.

To confirm the differential expression of key effector molecules between stimulation with cognate Ags versus PMA/I, we measured IL2, IL10, TNFɑ, and IFNγ protein concentrations in the culture supernatant after 18 hours of stimulation (Figures 3F and 3G, and Figures S3D and S3E). The levels of secreted IL2 and TNFɑ were significantly higher upon stimulation with PMA/I relative to DCs pulsed with cognate Ag (Figures 3F and S3D). IFNγ concentrations were similar between PMA/I and cognate Ags, while IL10, known to mediate TCR-signaling negative feedback, was significantly increased only by TCR engagement with cognate Ags, confirming RNA expression data.

In summary, the transcriptome analysis indicated that exposure to cognate Ags engaged pathways critical for viral reactivation. However, there were clear differences in patterns of gene expression induced by cognate antigen compared to those induced by polyclonal stimuli, with the largest differences observed with PMA/I. Treatment with PMA/I led to global changes in T cell transcription, including greater magnitude and breadth of transcription from NF-κB and NFAT-regulated genes. These results highlight the importance of directly assessing the ability of cognate antigens to induce HIV-1 gene expression.

### Re-encounter with cognate Ag induces HIV-1 expression

To study induction of HIV-1 expression by different stimuli, we exposed the enriched, expanded populations of Ag-responded cells to different restimulation conditions and sorted CD154^+^CD69^+^ responding cells for nucleic acid extraction (workflow depicted in Figure 1A). When experimental conditions lead to insufficient upregulation of activation markers (NoTx, NoAg, and KLH), we sorted total live CD4^+^ T cells (see gating strategy Figure S2E). To assess HIV-1 expression, we quantified cell-associated polyadenylated (polyA) HIV-1 RNA transcripts, which include both unspliced and spliced HIV-1 mRNAs that have been completely transcribed and polyadenylated^60–62^. The data were normalized by calculating cell equivalents from corresponding measurements of the host gene RPP30 in genomic DNA^49,60^.

Basal HIV-1 expression in the untreated condition (NoTx) varied greatly across participants (Figures 4A and Figure S4A). The presence of measurable HIV-1 RNA in untreated cells (NoTx) is consistent with the fact that some cells may not have returned to a completely quiescent state after the initial expansion, as indicated by low level expression of CD69 activation marker captured by our gating strategy (Figure S2E). Non-specific stimulation with PMA/I increased HIV-1 RNA expression in all 10 participants. Additionally, stimulation with PMA/I caused higher upregulation of HIV-1 RNA compared to anti-CD3/CD28 in 7 out of 10 participants. Most importantly, physiological stimulation with cognate Ag processed and presented by autologous DCs caused a statistically significant increase in HIV-1 RNA in 9 out of 10 study participants compared to NoTx, NoAg and KLH controls (Figure 4A). The median fold changes in HIV-1 RNA relative to NoTx upon stimulation with PMA/I and CD3/CD28 were 6.47 and 4.77, respectively, whereas upon re-encounter with cognate Ag the median fold change was 3.98 (Figure 4B). Overall, re-encounter with cognate Ag induced increased HIV-1 expression, but this increase showed inter-participant variability (fold change to NoTx range, 0.83-58.4; p=0.017). We observed the highest increase in HIV-1 RNA relative to NoTx for participant P3 (46.6-fold change; p<0.0001) and P4 (58.4-fold change; p<0.0001), whereas there was no significant fold change for participant P9 (0.83; p=0.86) (Figure 4B). Increase in HIV-1 RNA was specific to encounter with cognate Ag and statistically significant compared to NoAg and KLH (median fold change to NoAg 3.64, p=0.0004, Figure 4C). Furthermore, our results showed a significant increase in HIV-1 RNA/HIV-1 DNA ratio in PMA/I, anti-CD3/CD28 and cognate Ag stimulation compared to NoTx (median values 4.43, 5.32 and 3.84, respectively) (Figure 4D). Additionally, to understand if pharmacological LRAs can cause HIV-1 expression in our system, we measured HIV-1 RNA from sorted CD4^+^ T cells upon stimulation with bryostatin. Although bryostatin caused an increase in HIV-1 RNA in 5 out of 10 individuals, this was not statistically significant (Figure S4B). Our results are consistent with previous studies which showed single LRA treatment to be inefficient in reversing latent HIV-1 from PWH^63,64^.

**Figure 4.**
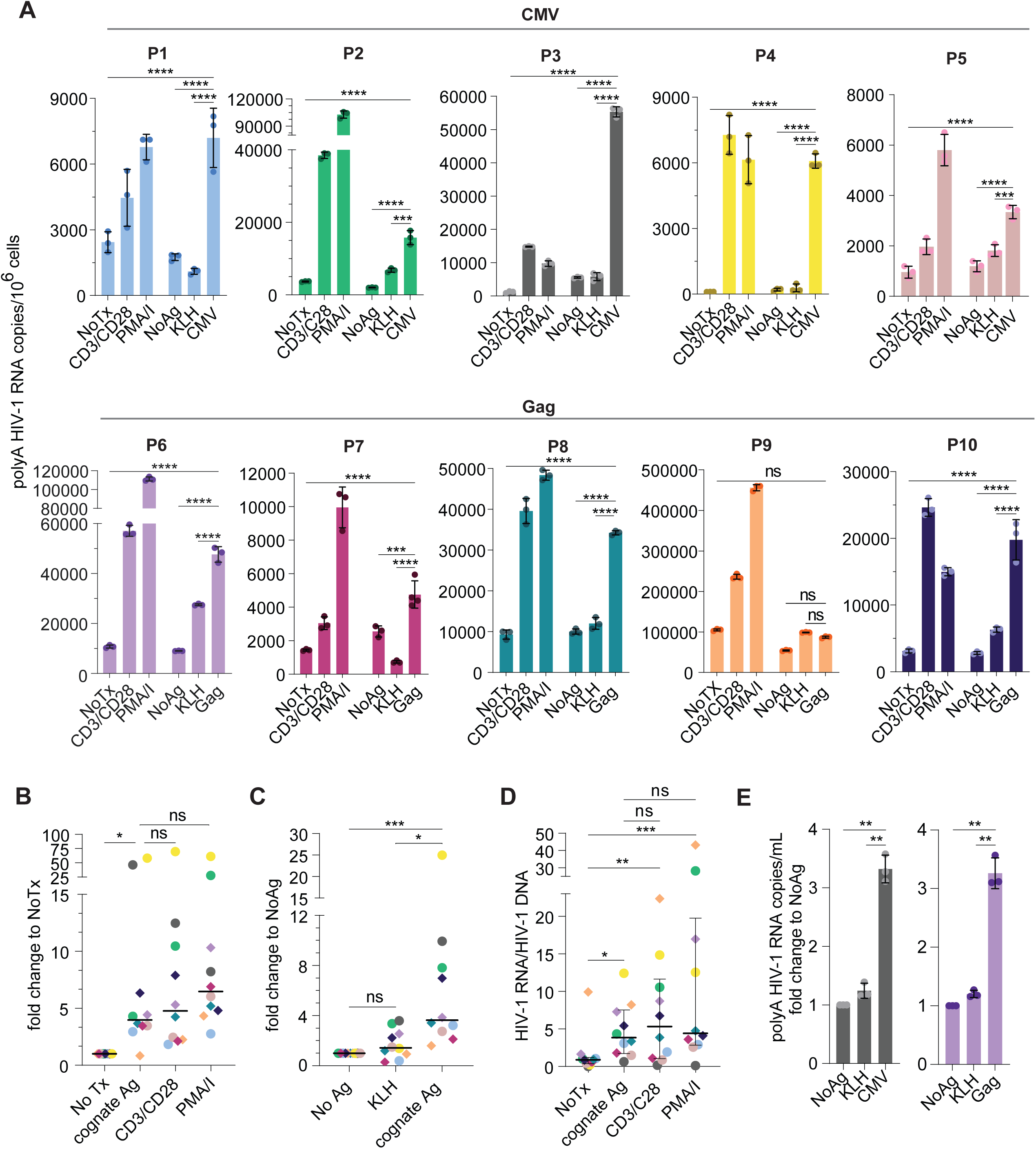
Encounter with cognate Ag causes significant increase in HIV-1 expression. (**A**) Quantification of HIV-1 polyadenylated RNA in CMV-responding cells (top panel; participants P1-P5) and Gag-responding cells (bottom panel; participants P6-10) across different treatments conditions. Data reported as HIV-1 polyA RNA copies per million cell equivalents; error bars indicate standard deviation; circles indicate technical replicates of ddPCR. Significance was determined by one-way ANOVA followed by Šídák’s test for multiple comparisons. (**B**) HIV-1 RNA expressed as fold change relative to NoTx (**C**) HIV-1 RNA expressed as fold change relative to NoAg. (**B-C**) Each colored circle represents the mean HIV-1 RNA value for a single participant. Circle colors match participants’ IDs. Statistical significance between conditions was determined by one-way ANOVA followed by Dunn’s test for multiple comparisons. (**D**) HIV-1 RNA induction expressed as HIV-1 polyA RNA and HIV-1 DNA ratio for NoTx, PMA/I and cognate Ag. HIV-1 DNA is used as a total number of proviruses calculated by IPDA (see Figure 1D). Single dot represents an individual study participant. Horizontal bars represent median with interquartile range. Significance was determined by one-way ANOVA followed by Dunn’s test for multiple comparisons. (**E**) Quantification of virion-associated HIV-1 RNA in supernatant collected after 18-hour coculture of expanded Ag-responding cells and autologous DCs loaded with NoAg, irrelevant Ag-KLH or cognate Ag, expressed as fold change to NoAg. Statistical significance between conditions was determined by one-way ANOVA. *P<0.05; **P<0.01, ***P<0.001, ****P<0.0001.

To understand the relationship between Ag-mediated T cell activation and latency reversal, it is important to consider whether the efficiency of Ag processing and presentation by DCs affects the strength of T cell responses^65^. Co-culturing CD4^+^ T cells with DCs loaded with protein Ag resulted in induction of HIV-1 RNA in 3 of 5 donors. The level of induction was lower than that observed with peptide-pulsed DCs, although this decrease was not statistically significant (Figure S4C). Our results demonstrate that presentation of epitopes derived from natural processing of the Gag P55 protein antigen by autologous DCs can reverse latency in Gag-responding CD4^+^ T cells.

We then investigated whether levels of HIV-1 RNA correlated to T cell activation and frequency of infected cells. The percentage of CD69^+^CD154^+^ cells did not correlate with HIV-1 RNA levels normalized by infected cell frequency (HIV-1 RNA/HIV-1 DNA) in either untreated cells (NoTx) or upon stimulation with PMA/I or cognate Ag (Figure S4D), likely due to the similar levels of CD69 and CD154 expression among participants in each condition. In addition, although T cell activation is required for latency reversal, the two processes are, to some degree, uncoupled^32,51,66^. As expected, we observed a significant correlation between HIV-1 RNA and the frequency of total proviruses (as calculated by IPDA, see Figure 1D) with both PMA/I and cognate Ag stimulation (Figure S4E).

To determine whether stimulation with cognate Ag would also result in viral particle production, we quantified HIV-1 RNA from viral particles in the supernatant from the co-culture experiments described above (Figure 4E). We analyzed samples from five participants, but only two had detectable HIV-1 RNA in supernatant, reflecting that most proviruses contributing to cell-associated RNA are defective^67,68^. In P3 and P6, we observed a significant increase in supernatant HIV-1 RNA copies upon stimulation with cognate CMV and Gag, respectively, relative to culture with DCs left unpulsed or pulsed with KLH (Figure 4E). Upon sequencing of HIV-1 RNA by limiting-dilution PCR, we identified a single variant in P3, matching the 5’-Leader defective provirus integrated outside of the *DELEC1* gene. Previous work from this participant demonstrated that this provirus is integrated in a dominant CMV-reactive clone (Figure S4F)^20^. In P6 we also detected a single variant, which matched one of the two proviruses contributing to cell-associated RNA (Figure S4G).

Taken together, our data demonstrate that a significant increase in HIV-1 RNA expression occurs upon stimulation with cognate Ag presented in a physiologic manner by autologous DCs. The median increase in HIV-1 RNA was lower with cognate Ag than with PMA/I. Furthermore, the magnitude of T cell activation as determined by upregulation of CD154 and CD69 among Ag-responding CD4^+^ T cells did not predict the levels of HIV-1 RNA expression. Finally, in the expanded pools containing proviruses capable of producing viral particles, we showed an increased release of specific viral variants upon stimulation with cognate Ag, suggesting that immune stimulation can contribute to the spontaneous transcriptional activity of the reservoir observed in vivo^69,70^.

### Cognate antigens and PMA/I induce comparable populations of infected cells

To further characterize HIV-1 expression induced upon re-encounter with cognate Ag or stimulation with PMA/I, we performed limiting dilution amplification and sequencing of cell-associated HIV-1 RNA from the sorted CD154^+^CD69^+^CD4^+^ T cells following restimulation to investigate whether similar populations of infected cells were induced to express viral RNA following polyclonal or Ag-specific stimulation. We obtained a total of 373 sequences from 6 participants, with an average of 62 sequences/participant (Figures 5A and Figure S5). We hypothesized that if PMA/I induced latency reversal for a larger proportion of proviruses, we would observe a higher number of distinct viral variants, including some than would not be found upon stimulation with CMV or Gag. With both forms of stimulation, the majority of the HIV-1 RNA sequences were dominated by a single or a few variants (Figure 5A and 5B). In all cases, the most abundant variants were detected in both stimulation with PMA/I and with cognate Ag, likely representing highly expanded infected clones and/or high RNA-producing cells (Figure 5B and 5C). We observed no difference between the proportions of identical sequences upon stimulation with PMA/I or cognate Ag (Figure 5D, median values 0.92 and 0.88, respectively). In contrast, a heterogeneous set of HIV-1 RNA sequences were recovered following stimulation of Gag-reactive cells from one participant, P8 (Figure S5A), suggesting the presence of a broad population of HIV-1-infected, Gag-reactive clonotypes. However, all of the most frequently recovered RNA variants were seen with both stimulations, and only rare RNA variants were unique to either PMA/I or cognate Ag stimulation. To test whether HIV-1 RNA sequences obtained with two types of stimulations were equally skewed by dominant variants, we calculated the Gini coefficient, a measure of distribution previously used to estimate oligoclonality of infected cells^71^. We observed no significant difference in Gini index between stimulation with PMA/I and cognate Ag (Figure 5E) and the median percentage of sequences induced both with PMA/I and cognate Ag was 86.7% (Figure 5C). Importantly, the enrichment and expansion of Ag-responding cells allowed us to test different stimuli with aliquots of the same cell population with comparable infection frequency and composition of proviruses, something which was not possible in previous approaches^42,43^. Our results show that in enriched populations of Ag-reactive cells, the same proviruses can be induced by T cell activation with both cognate Ag and PMA/I.

**Figure 5.**
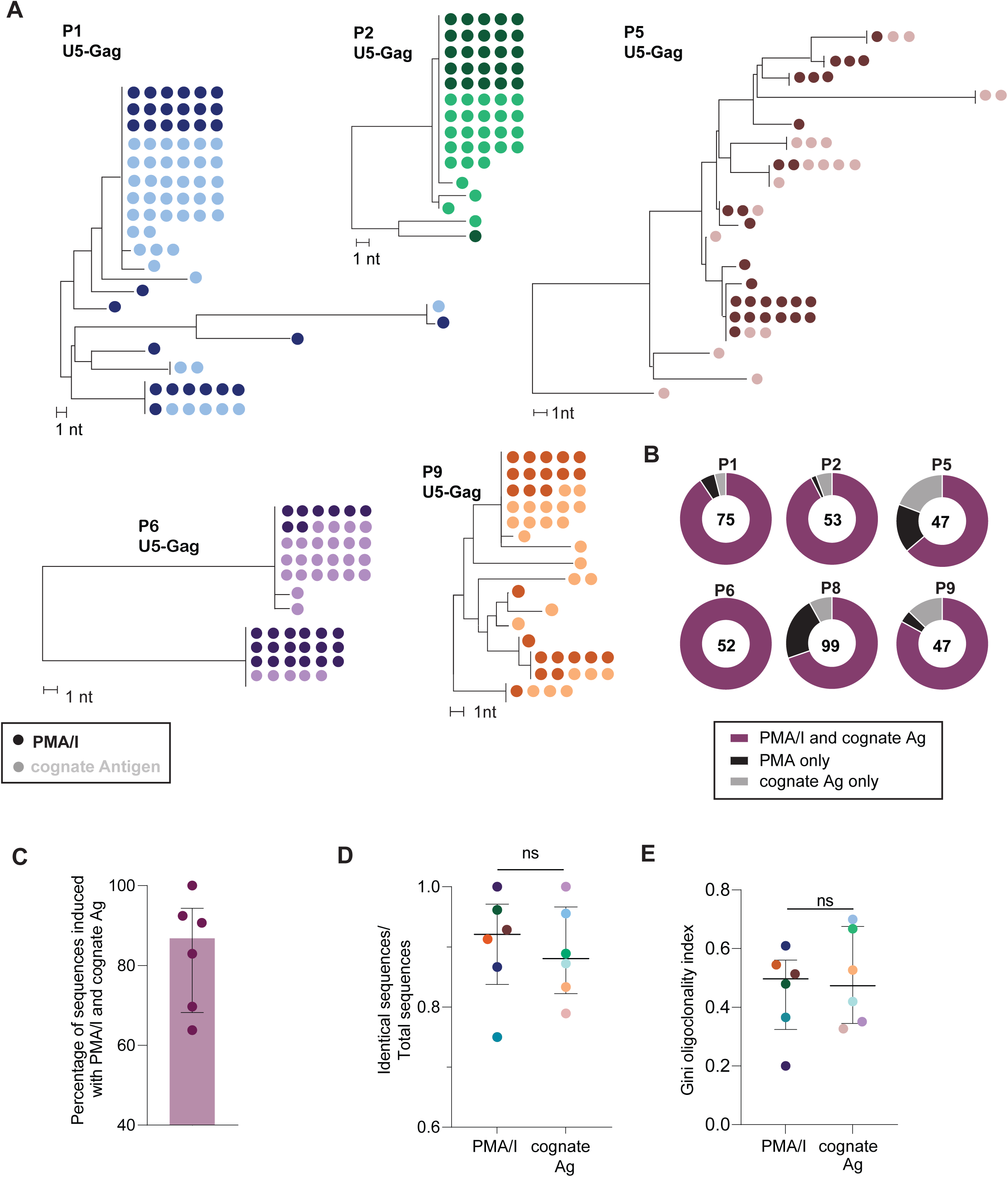
Stimulation with PMA/I and cognate Ag induce comparable populations of proviruses. (**A**) Representative neighbor-joining phylogenetic trees of cell-associated HIV-1 RNA u5-gag sequences from 5 study participants. Sequences recovered from PMA/I treatment are represented in dark shade, whereas those recovered after stimulation with cognate Ag are represented in light shade of corresponding color. A branch distance of 1 nucleotide (1nt) is shown on the tree scale. (**B-C**) Percentage of sequences found in both by PMA/I and cognate Ag for six study participants. The number of total recovered sequences is indicated in the center of the respective pie chart. (**D**) Frequency of identical sequences within sorted populations of either stimulation treatment for 6 study participants. (**E**) Gini oligoclonality index for 6 study participants. Horizontal bars represent median with interquartile range. (**D-E**) Horizontal bars represent median with interquartile range. Statistical significance was determined by paired t-test.

### Limiting-dilution analysis of HIV-1 RNA^+^ cells supports low proviral inducibility regardless of stimulation

Even though measuring cell associated HIV-1 RNA in bulk cultures of stimulated cells is a standard method to study latency reversal^64,72^, it does not provide information about differences in the induction of HIV-1 RNA at the single cell level. Indeed, the observed increases in HIV-1 RNA upon PMA/I or cognate Ag could be the result of modest transcription from many cells, or high-level RNA production from a small number of cells. Therefore, we optimized an assay to measure cell-associated HIV-1 RNA at the single HIV-1-infected cell level based on magnetic bead RNA isolation and cDNA synthesis^73^. In brief, small pools of cells were sorted into individual wells of 96-well plates, so that each well contained either zero or one HIV-1 RNA positive cell. To each well, we added 5000 PBMCs from a healthy donor to improve RNA recovery and spiked-in synthetic RNA as an internal control for RNA recovery. We then carried out RNA isolation, genomic DNA digestion, cDNA synthesis, and digital PCR to measure the number of HIV-1 polyadenylated RNA and internal control RNA copies per well (Figures 6A, Figure S6A and S6B, and methods). The assay sensitivity allows us to avoid pre-amplification, which could complicate the quantification of HIV-1 RNA copies within each cell^74^. We validated this method by quantifying commercially available HIV-1 RNA standards in 2-fold dilutions and confirmed the linearity of the assay, a 60% recovery of input HIV-1 RNA, and a 95% likelihood of detecting cells with at least 18 HIV-1 RNA copies (Figure S6C). The method is also specific, showed no evidence of inter-well cross-contamination between HIV-1 positive and negative cells, and allows the censoring of wells with poor RNA recovery using the internal RNA control (Figures S6D and S6E). To confirm this approach could detect changes in HIV-1 expression upon stimulation at the single infected-cell level, we first used well-characterized T cell lines that carry an HIV-1 provirus, ACH2 and JLat^75,76^. We assayed 45 singly-sorted ACH2 live cells that were cultured for 24 hours with PMA/I or left untreated, and observed a significant change in the average number of HIV-1 RNA copies/cell (15-fold increase, p<0.0001, Figure 6B). Additionally, to confirm we could assess biologically relevant changes in HIV-1 expression, we analyzed individual unstimulated JLat10.6 cells, which contain a GFP reporter provirus, sorted based on low, intermediate, or high GFP intensity (28 cells for each group, Figure 6C). We detected a significantly higher percentage of HIV-1 RNA^+^ cells among those with high GFP intensity (Figure 6D) (96%, versus 35% and 3.5% for intermediate and low GFP expression, respectively, p<0.001), which also contained significantly higher copies of HIV-1 RNA per cell (p<0.0001).

**Figure 6.**
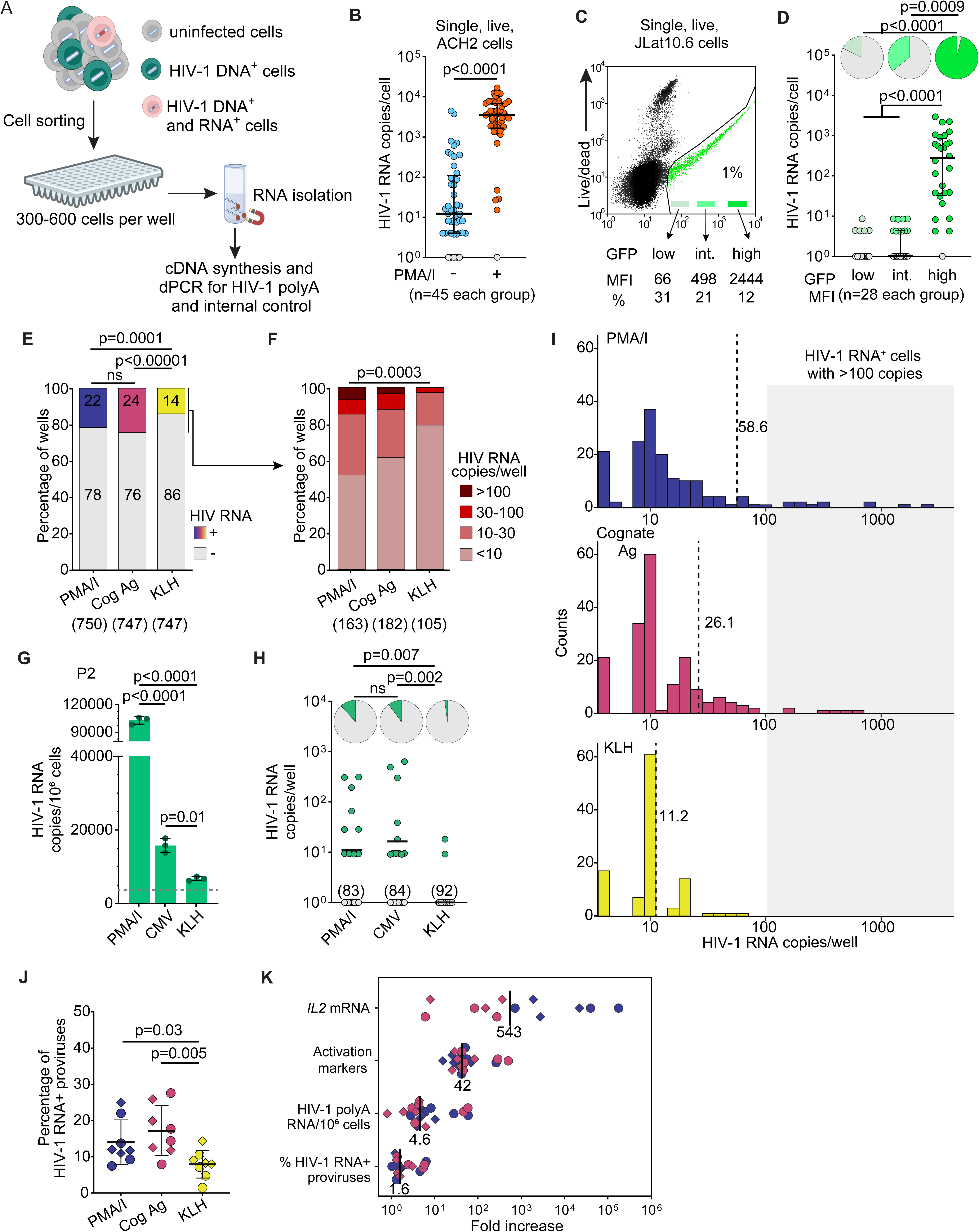
Quantification of HIV-1 RNA at single-infected cell resolution. (**A**) Experimental design; cells are sorted into 96-well plates so that each well contains only one or no HIV-1 RNA positive cells, either by single cell sorting or limiting dilution; a synthetic RNA spike-in is used to control for RNA recovery. (**B**) Individual ACH2 cells were analyzed after 18 hours of culture with and without PMA/I (orange and blue symbols, respectively); gray symbols indicate values below the limit of detection; error bars indicate median and interquartile range; the difference between the two groups was assessed by Mann-Whitney t-test. (**C**) Flow cytometry plot of unstimulated JLat 10.6 cells, showing sorting logic based on GFP fluorescence intensity. (**D**) Analysis of individual JLat 10.6 cells sorted based on low, intermediate, or high GFP expression; gray symbols indicate values below the limit of detection; error bars indicate median and interquartile range; the difference between the three groups was assessed by Kruskal-Wallis test; pie charts at the top represent the percentage of HIV-1 RNA^+^ cells; the difference between the three groups was assessed by Fisher’s exact test. (**E**) Percentage of all wells positive for HIV-1 RNA grouped by treatment with PMA/I, cognate Ag or irrelevant KLH; numbers in parenthesis indicate the total wells analyzed. (**F**) Percentage of wells grouped by HIV-1 RNA copies; the difference between the three groups was assessed by Fisher’s exact test. (**G**) HIV-1 RNA quantification in bulk cells, as in Figure 4A, for participant P2; the dashed horizontal bars indicate the mean value for cells without treatment (NoTx). (**H**) Limiting dilution viral quantification assay for participant P2; black horizontal bars indicate mean values; gray symbols indicate values below the limit of detection (9 copies/well) and numbers in parenthesis indicate the number of negative wells; pie charts at the top indicate the percentage of HIV-1 RNA^+^ wells; the difference between the three groups was assessed by Fisher’s exact test. (**I**) Histograms showing the distribution of HIV-1 RNA copies/well; dashed lines indicate the mean values. (**J**) Percentage of induced proviruses calculated as the ratio between the number of positive HIV-1 RNA wells and total number of proviruses seeded in each plate. Each dot represents one plate from each condition and study participant. Horizontal lines represent median value. Significance was determined one-way ANOVA. Estimate of total cells and proviruses screened in each plate; horizontal bars indicate mean values. (**K**) Comparison of fold changes in IL2 mRNA expression, T cell activation-induced markers, and HIV-1 expression in bulk or at the single-infected cell level, relative to negative controls; each dot represent experiments from one participant; blue and magenta indicate stimulation with PMA/I and cognate antigens, respectively; circles and diamonds indicate experiments with CMV and Gag-reactive cells; black bars and numbers indicate median values.

We then applied this approach to the same ex vivo expanded CD4^+^ T cells from which we quantified HIV-1 RNA in bulk cultures after stimulation with PMA/I or DCs loaded with cognate Ag. Expecting that only a small fraction of cells would be infected and transcriptionally active, we sorted small pools of CD154^+^CD69^+^ cells into microtiter wells (300 to 600 cells per well) so that less than 30% of wells would contain an HIV-1 RNA^+^ cell. Pools with the same number of total CD4^+^ cells were sorted for samples restimulated with DCs loaded with KLH. We studied 24 samples from 8 participants, with an overall RNA recovery of 62% (standard deviation ±8%) based on the internal control, and a total of 2244 individual wells assayed (Figure S6F). As expected, most wells were HIV-1 RNA negative (80%), but we detected a significantly higher percentage of HIV-1 RNA^+^ wells from samples treated with PMA/I or cognate Ag compared to KLH (Figure 6E). Overall, the mean values of HIV-1 RNA copies/well were 12.06, 6.36, and 1.56 for cells treated with PMA/I, cognate Ag, and KLH, respectively (p=0.012). Moreover, we detected more cells with higher copies of HIV-1 RNA from PMA/-I and cognate Ag-stimulated cells compared to those treated with KLH (p=0.0003), which had a very few cells with >30 HIV-1 RNA copies (15% and 12%, versus 3%, p=0.01, Figure 6F). Indeed, the direct comparison of HIV-1 RNA quantification from cells in bulk versus cell at limiting dilution revealed that differences in HIV-1 gene expression were mostly driven by rare cells with higher copies of HIV-1 RNA (>100 copies/cell). Figures 6G and 6H show a representative example from participant P2, in which we detected high RNA-producing cells only with PMA/I and DCs pulsed with CMV Ag. Conversely, in participant P9 only PMA/I induced high RNA-producing cells, in agreement with the HIV-1 RNA quantification in bulk (Figures S6G and S6H). Overall, we detected rare high-producing HIV-1 RNA^+^ cells (>100 copies) in 5 out of 8 participants (Figure 6I), corresponding to only 3.8% of all HIV-1 RNA^+^ cells. Because these cells are extremely rare among all CD4^+^ T cells, bulk measurements cannot provide this single-infected cell resolution, but they can be better suited to detect differences among samples. In the pooled analysis of HIV-1 RNA^+^ wells from all 8 participants, although PMA/I had the highest occurrence of high RNA producing-cells and mean copies/well (58.6±256), the difference in HIV-1 gene expression among the three groups did not reach statistical significance (p=0.14, Figure 6I), likely reflecting the low proviral inducibility even among activated (CD69^+^CD154^+^) cells. Finally, to provide an estimate of HIV-1 RNA^+^ cells among those infected, we calculated the percentage of positive wells for HIV-1 RNA and the number of total proviruses spread across the plate (Figures 6J and S6I). We estimated that only 15% (±6) of proviruses in cells activated with PMA/I or cognate Ag were positive for HIV-1 RNA (averages of 8 participants were 14% and 17%, respectively, Figure 6J). Albeit low, this percentage was significantly higher than CD4^+^ T cells exposed to DCs pulsed with KLH (average 7% ±3, p=0.005). Even though exposure to cognate Ag and stimulation with PMA/I resulted in robust CD4^+^ T cell activation (as indicated by upregulation of IL2 mRNA expression and CD69 and CD154 activation markers), we observed only a modest increase in HIV-1 expression with both bulk and single cell measurements (median fold change 4.6 and 1.6, respectively) (Figure 6K).

Overall, the use of a limiting dilution assay helped us distinguish between low (1-10 copies/cell), moderate (10-100 copies/cell) and high (more than 100 copies/cell) HIV-1 RNA producing cells. This striking variation in viral gene expression among infected cells would have been missed with bulk RNA assays or less sensitive single-cell approaches^25^. Additionally, stimulation with cognate Ag or PMA/I similarly induced high RNA-producing cells, despite the low inducibility of proviruses persisting on long-term ART, as shown in earlier studies.^32,77^

## Discussion

Even in PWH who have maintained optimal adherence to ART, the latent reservoir persists in resting CD4^+^ T cells, including cells that have undergone clonal expansion. Indeed, previous work supports a major role for antigen-driven clonal expansion in promoting HIV-1 persistence^20,24^, and most of the HIV-1 reservoir is made of expanded CD4^+^ T cell clones^15,16,19,21,78^. These clones fluctuate in size over time, with temporal dynamics resembling the expansion and contraction of immune responses upon Ag exposure^79^. Previous work characterized HIV-1-infected, CMV-responding cells in PWH and showed that these populations were dominated by large clones of infected cells^24^. Understanding which immunological factors dictate the capacity of infected clones to proliferate and survive is paramount for the development of novel curative strategies. However, there is limited understanding of the linkage between Ag-mediated T cell activation, infected cell proliferation, and the induction of HIV-1 gene expression. Although it is clear that polyclonal T cell activation by mitogens, anti-CD3/CD28, or PMA/I can reverse latency, there has been limited proof that, in purified populations of Ag-responding cells, exposure to Ag presented in a physiologic manner by autologous DC can reverse latency. Studying Ag-mediated latency reversal is challenging because of the low frequency of HIV-1 infected CD4^+^ T cells specific for any particular antigen. Here, we implemented a novel approach to: 1) expand the population of Ag-responding CD4^+^ T cells to study the infection rate and proviral composition among these cells; 2) determine whether encounter with cognate Ag presented in a physiologic manner by autologous DCs can cause induction of HIV-1 expression; 3) compare the transcriptional activation and HIV-1 expression induced by cognate Ag versus non-specific T cell activators in a controlled setting.

Dendritic cells are professional antigen presenting cells (APCs) critical for the initiation of adaptive immune responses. Antigen-specific T cell receptors (TCRs) are constantly performing dynamic surveillance for cognate peptides presented on the surfaces of APCs^52,80,81^. Encounter with cognate-pMHC complexes and the formation of an immunological synapse are highly regulated multistage processes which dictate physiologic T cell activation^52,80^,. A previous study emphasized the importance of cross-talk between lymphocytes and DCs, including the CD40/CD40L (CD154) axis, in the context of HIV-1 reactivation^43^. However, interpretation of the results was complicated by the lack of antiretroviral drugs needed to distinguish latency reversal from viral replication^43^. In another study, stimulation of CD8-depleted PBMCs with pools of peptides derived from HIV-1 and other common pathogens (CMV, EBV, Candida, flu, tetanus) led to inconsistent latency reversal across patients. These findings suggested that infected cells are not commonly enriched in any particular population of Ag-specific memory cells. However, analysis was complicated by the rarity and heterogeneity of proviral populations across cells susceptible to different stimulations, which prevented the comparison between peptide pools and non-specific, positive controls (anti-CD3/CD28)^42^. Here, we used CMV and HIV-1 Gag as frequently encountered Ags and isolated CMV- and Gag-responding cells as previously described^36,45^. We then expanded these cells to quantify the frequency of genetically intact and defective proviruses in the expanded populations by IPDA. Previous studies suggested that a subset of CD4^+^ T cells specific to CMV are less frequently infected – and spared during untreated infection – relative to cells with other specificities due to high autocrine production of MIP1-β^82^. Conversely HIV-1-reactive cells may be preferentially infected during acute infection and viral rebound^83,84^. Despite the limited number of participants, we did not find a significant difference in infection frequency between CMV- and HIV-1-responding cells expanded ex vivo. Although we found enrichment for total and intact HIV DNA within Ag-reactive cells in some participants, IPDA data showed striking heterogeneity across participants suggesting disparate distribution of proviruses among CD4^+^ T cells with distinct antigenic specificities. This notion that the latent reservoir does not reside in CD4^+^ T cells of only one subset or with recurring antigen specificities was suggested in earlier work^20,24,32,42^. Indeed, our data showed that HIV-1 can persist in memory CD4^+^ T cells of diverse Ag-specificity. Despite the limited range of antigens tested in this study, our results suggest that the Ag-specificity of HIV-1 infected cells differs from person to person, and that HIV-1 cure strategies that would reduce the antigenic stimulation of specific infected clones (e.g. reducing exposure to CMV Ags with antivirals) would have little effect on the latent reservoir globally in most individuals.

Upon restimulation of the expanded pools of Ag-reactive cells, we observed the highest upregulation of CD154 and CD69 in cognate Ag-dependent manner in all participants, indicating that the enrichment and expansion protocol gave a highly specific populations of CD4^+^ T cells responsive to the relevant Ag. Stimulation of CD4^+^ T cells with anti-CD3/CD28 and PMA/I also led to significant increase in activation-induced markers. However, transcriptome analysis demonstrated that stimulation with PMA/I, which bypasses the early regulatory steps in TCR signaling, results in significantly higher transcriptional activation and expression of effector molecules (IL2, TNFɑ, etc.), whose genes are cooperatively regulated by NF-κB and NFAT^85,86^. Indeed, it is likely that because of this integration of kinase and calcium signaling, PMA/I is one of the strongest HIV-1 latency reversing strategies ex vivo^63,87^.

Using autologous DCs loaded with either CMV or HIV-1 Gag antigens, we demonstrated significant increases of HIV-1 RNA expression in sorted CD154^+^CD69^+^CD4^+^ T cells relative to NoTx and NoAg controls in cells from 9 out of 10 study participants. These results provide a direct demonstration that encounter with cognate Ag in a physiologic context can increase expression of HIV-1 genes, a finding with direct relevance for the critical question of what causes HIV-1 rebound upon treatment interruption.

Even though T cell activation is a prerequisite for latency reversal^88^, our data suggest that there is some discordance between Ag-driven T cell responses and latency reversal. This observation is consistent with studies demonstrating that for some proviruses latency reversal was achieved only after multiple rounds of T cell activation^16,32^, and that other factors, such as proviral location, can further reduce viral expression despite T cell activation and proliferation^66^. In addition, we showed that higher infection frequency among Ag-responding cells resulted in higher HIV-1 expression upon cognate Ag and PMA/I stimulation.

Although the increase in HIV-1 RNA was the highest with PMA/I in 6 out of 10 individuals, the median fold changes relative to NoTx upon PMA/I or Ag stimulation were not statistically different (Figure 4B). In addition, there were no observed differences in the populations of proviruses induced by these stimuli. Our sequencing data confirmed that stimulation with both PMA/I and cognate Ag frequently induced identical HIV-1 RNA variants. Overall, our results show that at the level of individual infected cells, physiologic stimulation with cognate Ag can induce HIV-1 gene expression to an extent comparable to that seen with polyclonal activators like PMA/I despite the observed differences in the transcriptional changes induced by these stimuli.

To examine differences in HIV-1 RNA induction at the single infected-cell level, we optimized a sensitive assay that allows to quantify polyadenylated HIV-1 RNA from either individual cells or small pools at limiting dilution for HIV-1 expressing cells. A similar approach was used in a previous study to quantify viral RNA transcripts from single proviruses in individuals on ART receiving a short treatment with the LRA romidespsin, which triggered a small number of memory CD4^+^ T cells to become only marginally transcriptionally active^89^. Conversely, we observed that stimulation of the Ag-responding expanded pools with either PMA/I or cognate Ag led to the induction of rare cells with higher HIV-1 expression levels, which contributed to the fold-increase in viral transcripts measured from CD4^+^ T cells in bulk. Despite the detection of these high HIV-1-expressing cells, our results point towards the overall low-inducibility of proviruses that persist in individuals on long-term ART. Indeed, we estimate that after activation by TCR signaling with different stimuli, only about 15% of infected cells expressed detectable levels of polyadenylated HIV-1 transcripts (≥10 copies/cell). This observation further supports a model in which the HIV-1 reservoir persists in part due to a higher threshold for latency reversal^60^ relative to the transcriptional activation required for T cell responses. This allows most infected cells to proliferate and deploy effector functions while avoiding immune recognition. Our findings also highlight the need to develop new latency reversing agents that target HIV-1-specific regulators of RNA transcription rather than broad T cell activation, as recently reported^63,90^.

Previously, a direct comparison of HIV-1 gene expression upon polyclonal and specific immune stimuli was impossible, due to the much smaller proportion of cells reactive to a single antigen, and the difference in the fraction of total and intact proviruses in responding populations. Our experimental design “leveled the playing field”, allowing us to directly compare proviral inducibility in response to *i)* physiologic antigen presentation by DCs versus polyclonal stimulation based on *ii)* CD3/CD28 engagement or *iii)* PKC/Calcium signaling. These stimuli led to different patterns of transcriptional activation but ultimately caused similar levels of T cell activation with detectable but restricted levels of HIV-1 gene expression. This result has important implications for recent studies on reservoir persistence in long-term survivors who have been on ART for more than 20 years. Integration site analyses paired to HIV-1 genome sequencing indicated that intact proviruses are progressively enriched in heterochromatic regions linked to deeper latency^91,92^, yet our group has recently found many readily inducible infectious proviruses in such individuals^12^. The latter finding was based on the QVOA, which relies on latency reversal with the mitogen phytohemagglutinin (PHA), raising the question of whether polyclonal activation with mitogens induced proviruses that would not be induced in vivo. Here, we extend the results from a recent case report^66^ and show that Ag presented in a physiologic manner by autologous APCs can reverse latency and cause viral particle production, further supporting that even proviruses that have been selected during long-term ART can be reactivated and could contribute to viral rebound if ART is interrupted in the absence of exceptional immune control.

In conclusion, this study shows how one of the fundamental functions of CD4^+^ T cells – the constant scanning and recognition of specific antigens – can affect the spontaneous transcriptional activity of the HIV-1 reservoir during ART. Although HIV-1 persistence is maintained through latency, residual viral gene expression driven by encounter with Ag may sustain anti-HIV-1 adaptive immunity^69^, contribute to immune activation^93^, and give rise to residual viremia that can complicate ART management^94^. Most importantly, our findings suggest that antigen recognition may affect the earliest events leading to viral rebound when ART is stopped. Therefore, future studies are needed to develop interventions that would eliminate virus-producing cells and prevent viral dissemination, since antigenic stimulation could eventually overcome even deep proviral latency.

## Limitations of the study

The major limitations of the study were the striking heterogeneity in proviral frequency and composition within antigen-reactive cells across participants, the inability to determine the intactness of the induced proviruses that contributed to both bulk and limiting-dilution RNA measurements, and the lack of integration site data to study the impact of genomic location on proviral inducibility. In future studies, our approach could be used to sort HIV-1-infected cells upon re-stimulation based on intracellular p24 staining^69,84^ or fluorescent in situ hybridization^69^, allowing in depth characterization of proviruses in the Ag-responding cells expanded ex vivo. Another technical challenge of this work was related to the use of a polyclonal population of Ag-responding cells stimulated with lysates or peptide pools. This approach prevented us from differentiating the contribution of TCR avidity, signaling strength and pMHC density to proviral inducibility. However, the identification and manipulation of individual infected clones with known specificity at the single epitope level is extremely challenging. An additional limitation of the study is that it focused on only 10 individuals, including only 3 females; hence, the generalizability of our conclusions to people of different ages, sexes, and ethnicities remains to be determined.

## Acknowledgements

We thank the study participants who volunteered to take part in this study and their families. We thank Joel Pomerantz, Joel N. Blanson, and Petra Bacher for discussions leading to this work, and Alicia Edwards for administrative support. This work was supported by the NIH NIAID Martin Delaney I4C (UM1AI164556), Beat-HIV (UM1AI126620), DARE (UM1AI164560), and PAVE (UM1AI164566) Collaboratories. This work was also supported by the Office of the NIH Director and National Institute of Dental & Craniofacial Research (DP5OD031834) (FRS), the Johns Hopkins University CFAR (P30AI094189) (FRS), the Robert I. Jacobs Fund of The Philadelphia Foundation (LJM), the Herbert Kean, M.D., Family Professorship (LJM), and by the Howard Hughes Medical Institute (RFS).

Figures 1A, 3A and 6A were created by using resources from biorender.com.

## Authors contribution

M.M, R.F.S., and F.R.S. conceptualized the study. M.M. and F.R.S. designed the experiments and performed analyses. M.M., F.D., N.L.B, F.W., and F.R.S. performed experiments. F.D. conducted the implementation and optimization of the limiting-dilution virus quantification assay. H.Z. conducted cell sorting experiments. J.R.W. and F.R.S. conducted RNA-seq analyses. J.L., R.H., K.M., P.T., K.L., L.J.M., and S.G.D., enrolled the study participants and gathered clinical data. M.M., J.D.S., R.F.S, and F.R.S wrote the manuscript with feedback from all authors.

## Methods

### Study participants

Characteristics of the study participants are provided in Supplemental Table S1. Participants were persons living with HIV-1 (PWH) on suppressive ART initiated during chronic infection, with 8 out of 10 study participants having undetectable plasma HIV-1 RNA levels (<20 copies/mL) for 8 or more years. Participant P6 experienced intermittent periods of detectible viremia with a median level of 75 copies/ml (range <20-300 copies/ml), despite optimal adherence and no drug resistance. Previous work from our group has shown that P6’s nonsuppressible viremia is due to expanded CD4^+^ T cells carrying multiple proviruses, some with 5’-L-defects^95^ (study participant ID P2^95^). Participant P9 experienced intermittent nonsuppresible viremia despite optimal adherence since 2015 (mean HIV-1 RNA 26 copies/mL, median <20copies/mL). Additional inclusion criteria were CD4^+^ T cell count above 400 cell/μL and positive CMV serology (participants P1-P5). Peripheral blood samples (up to 180 mL) were collected at the Johns Hopkins Bartlett Specialty Clinic and processed to isolate PBMCs by Ficoll gradient separation. For participants P4, P5, P9 and P10, leukapheresis was performed at University of Pennsylvania and University of California San Francisco.

### Enrichment of Antigen-responding cells

Antigen-responding cells were enriched using the previously described antigen-reactive T cell enrichment (ARTE) method^36,44^. In brief, between 9 and 4 x 10^7^ CD8-depleted PBMCs were resuspended in RPMI-1640 (GIBCO), supplemented with 10% (v/v) human AB-serum (Sigma Aldrich,, Germany) seeded into well culture plates and stimulated with antigens for 18 h in presence of 1 mg/ml of anti-CD40 blocking antibody (both Miltenyi Biotec, Germany) and costimulatory molecules (CD28/CD49d; BD Biosciences; 0.5 μg/mL) as well as a cocktail of antiretroviral drugs (10 nM dolutegravir, 10μM tenofovir, and 10μM emtricitabine) to prevent new infection events. Cells stimulated with either lysates of CMV-infected fibroblasts (Virusys, 10 μg/mL), overlapping Gag 15mer peptides (HIV-1 Gag peptide pool, JPT peptides, 1 μg/peptide/mL) or Gag P55 protein (NIH reagents: HIV-1 IIIB pr55 Gag Protein, ARP-3276 and HIV-1 NL4-3 pr55 Gag Recombinant Protein, ARP-13385; 2.5 μg/ml). To study Gag-responding cells, we used 15mer overlapping HIV-1 Gag peptides as well as whole protein to better simulate natural antigen processing and presentation^96^. After 18 hours, cells were washed and labeled with CD154-Biotin followed by anti-Biotin MicroBeads (CD154 MicroBead Kit; Miltenyi Biotec) and magnetically enriched by two sequential MS columns (Miltenyi Biotec). Frequencies of antigen-responding T cells were determined based on the total cell count of CD154^+^ T cells after enrichment, normalized to the total number of cells applied on the column.

### In vitro expansion, preparation of dendritic cells (DCs) and re-stimulation of antigen-responding T cells

The negative cell fraction resulting from the CD154 magnetic enrichment was irradiated and used as antigen-leaded, autologous feeders for cell expansion ex vivo. Purified CD154^+^ T cells were expanded in presence of 1:100 autologous antigen-loaded irradiated feeder cells in TexMACS medium (Miltenyi Biotec), supplemented with 5% (v/v) human AB-serum (GemCell), 200 U/ml IL-2 (R&D) and 100 IU/ml penicillin, 100 mg/ml streptomycin, 0.25 mg/ml amphotericin B (Antibiotic Antimycotic Solution, Sigma Aldrich) and antiretrovirals. During expansion for 2 weeks, the medium was replenished, and cells were split as needed. To generate monocyte-derived dendritic cells, we cultured autologous CD14^+^ monocytes isolated by positive selection (CD14 MicroBeads; Miltenyi Biotec) in X-Vivo15 medium (BioWhittaker/Lonza), supplemented with 1000 IU/ml GM-CSF and 400 IU/ml IL-4 (both Miltenyi Biotec) for 5 days. On the second day, CD14^+^ cells were stimulated with the same antigens used for the CD154^+^ cell enrichment. Before re-stimulation, expanded T cells were rested in RPMI-1640 + 5% human AB-serum for 2 days. Expanded cells were re-stimulated in a coculture ratio of 1:1 with antigen-loaded fastDCs in 24-well flat bottom plates for 18 hours.

### Flow cytometry

Cells were washed and incubated with FcgR Block (BD Pharmingen) at room temperature for 10 minutes, and then stained for 30 minutes on ice with an APC-labeled antibody against CD3 (BioLegend; clone UCHT1), phycoerythrinCy7–labeled (PE-Cy7–labeled) antibody against CD4 (BioLegend; clone RPA-T4), BV421-labeled antibody against CD154 (BioLegend; clone 24-31), FITC-labeled antibody against CD69 (BioLegend; clone FN50), and PE-Cy5–labeled antibodies against CD14 (Thermo Fisher Scientific; clone 61D3) and CD16 (BioLegend; clone 3G8). Dead cells were excluded using propidium iodide. Cells stained with single fluorophore-labeled antibodies, costimulation-only controls, and positive controls were used to set sorting gates. Cells were sorted using either the Beckman Coulter MoFlo Legacy or the XDP cell sorter. A representative gating strategy and sorting logic is provided in Supplemental Figure S2E. Data was analyzed using FlowJo v10 (Treestar, Ashland,OR, USA) software.

### Extraction of genomic DNA (gDNA) and intracellular RNA

Sorted cells were collected and immediately lysed for nucleic acid extraction. Isolation of gDNA and cell-associated RNA was performed based on previously published protocols^49^. For samples with more than 10^6^ cells, QIAamp DNA Mini Kit (QIAGEN) was used for DNA isolation. NanoDrop 2000 and Qubit 3 Broad Range (Thermo Fisher Scientific) were used to quantify gDNA concentrations.

### Intact proviral DNA assay (IPDA)

Expanded cells were rested for 2 days and aliquots of 3-5 x 10^6^ CD4^+^ T cells were collected to isolate gDNA. The IPDA was performed as previously described^49^. Because of sequence variation, a custom ψ-probe was designed for participant P2 (FAM-TGGCGTACTCACCAGG-MGBNFQ; Applied Biosystems), and a custom ψ-forward primer was designed for participant P2 (CAGGACTCGGCTTGCTGAGC)^20^. Copies of intact and defective proviruses were corrected by DNA shearing and normalized to cell equivalents based on RPP30^49^.

### HIV-1 RNA quantification

Intracellular RNA was subjected to complementary DNA (cDNA) synthesis using random hexamer and oligodT and the SuperScript III First-Strand Synthesis System (Thermo Fisher Scientific) in order to measure HIV-1 poly-adenylated RNA transcripts. cDNA was quantified using primers and probe previously published^62^ and subjected to digital droplet quantification using the BioRad system. Data was analyzed using QuantaSoft Analysis Pro. Data were normalized to copies per 10^6^ cells by calculating gDNA cell equivalents based on RPP30.

### Single-Genome-Sequencing (SGS)

Cell-associated RNA was used for reverse-transcription with a gene-specific primer in Gag (TGACATGCTGTCATCATYTCYTC). The cDNA was then subjected to end-point dilution and PCR-amplified, and single-genome sequences were obtained from an 1,100-bp region spanning from U5 to Gag (HXB2 positions 584-1841), as previously described^97^. Outer and inner PCRs were performed as previously published^97^. PCR products were sequenced by sanger sequencing. Table S2 summarizes the oligos used in this study.

### Bioinformatics analysis of HIV-1 sequences

Raw data from Sanger sequencing were analyzed in Geneious to resolve base call conflicts and eliminate sequences with poor quality or double peaks reflecting more than one HIV-1 variant per PCR reaction. Sequence contigs were aligned with ClustalW^98^. ElimDupes was used to identify and collapse identical sequences (https://www.hiv.lanl.gov/content/sequence/elimdupesv2/elimdupes.html). Neighbor-joining (NJ) trees were constructed in MEGA 7.0 with a subtype-specific HIV-1 consensus as the outgroup^99^. The phylogenetic structure was tested by bootstrap analysis (1000 replicates). To investigate differences between RNA variants, we calculated the Gini coefficient of inequality in RStudio with the ineq R package (https://cran.rproject.org/web/packages/ineq/index.html) and corrected for small samples. This measurement of sample dispersion provides an estimate of whether the RNA variants within a sample are evenly distributed in groups of different sequences (values approaching 0) or dominated by groups of identical variants (values approaching 1).

### Quantification of HIV RNA in limiting dilution experiments

The limiting-dilution assay was developed based on a high-throughput, sensitive, and cost-effective RNAseq protocol, named prime-seq, with minor modifications^73^. Briefly, individual or small pools of cells were sorted in 96-well plates (Eppendorf twin.tec PCR LoBind) in 50µl of lysis buffer, consisting in Buffer RLT Plus (Qiagen, Ref#1053393) and 1% β-mercaptoethanol, with each well containing a cushion of 5000 uninfected PBMCs. Plates were stored at -80 °C until processed. In restimulation experiments of antigen-reactive, expanded cells treated with PMA/I or Ag-loaded DCs, live CD154^+^CD69^+^CD4^+^ T cells were sorted in pools of 300-600 cells/well. As controls, total CD4^+^ T cells treated with KLH-loaded DCs were sorted in equal pools of cells/well. Upon thawing, cell lysates were treated with 20µg of Proteinase K and 1µl of 25mM EDTA and incubated at 50 °C for 15min. In addition, ∼8000 copies of a synthetic RNA internal control (IC) (see details below) were added to each well. The lysed samples were then treated with cleanup beads, prepared as previously described^73^, at a 1:3 ratio to maximize the recovery of RNA. After an incubation of 5min at room temperature, cleanup beads were immobilized with a magnetic plate holder and washed twice with 70% ethanol. Then, genomic DNA was digested on-beads using 1U of DNAse I (NEB M0303) at room temperature for 10 minutes. Following incubation, DNase was inactivated by adding 1µl of 100mM EDTA and incubation at 65 °C for 5 minutes. After two washes with 70% ethanol, beads were resuspended in 10µl of RT-solution, consisting of 30U of Maxima H Reverse Transcriptase (Thermo Fisher EP0753), 1µM of template-switching oligo and 1µM of HIV-tailed oligodT primers, to perform on-beads cDNA synthesis. The reaction was incubated at 42 °C for 1h and 30min. At the end of cDNA synthesis, 10µl of 5mM Tris-HCl were added to each well and the plate was vortexed thoroughly to ensure proper recover of cDNA from the beads. Finally, the samples were cleaned and concentrated in a final volume of 10µl of 5mM Tris-HCl using cleanup beads in a 1:1 ratio. Five µl of cDNA were quantified in a duplex digital PCR reaction targeting total HIV-1 polyadenylated RNA^60^ and the IC RNA. Digital PCR reactions were run using the QIAcuity One Digital PCR System (Qiagen) in a 96-well 8.5k partitions nanoplate, with an initial denaturation step of 95 °C for 2min, followed by 45 cycles each including 95 °C for 15min and 58°C for 1h. Samples from participant P6 were run in 24-well 26k partition nanoplates. Table S2 summarizes the oligos used in this study.

The analytical sensitivity of the assay was assessed by the quantification of 2-fold dilutions (input ranging from 250 to 2 copies) of a validated HIV RNA standard (HIV Reagent Program, ARP 3443). Twenty-four independent measurements were assayed for each point and the 95% and 50% hit rate were calculated by Probit regression analysis. The performance and the ability of the assay to detect RNA starting from a single HIV-1 RNA^+^ cell was tested on singly sorted ACH-2 cells with or without stimulation with PMA/I for 12h, and on unstimulated single J-Lat 10.6 cells sorted based on low, mid, and high GFP expression (See Supplemental Figure S6). ACH-2 and Jlat10.6 cells were obtained from the HIV Reagent Program (ARP-349 and ARP-9849, respectively).

### Synthetic RNA control

To properly account for RNA loss during the RNA extraction, we designed a control for recovery, consisting in a synthetic RNA, spiked-in in each well before the extraction process. Briefly, we designed a gblock (IDT DNA, Coralville, CA) of 2,728bp sharing homology with DENV NS3 gene, with the sequence being modified to optimize the gblock synthesis. To introduce the T7 promoter primers required for the in vitro transcription process, we used 100ng of the gblock in a PCR using 4µl of 10x Buffer, 2.5mM of MgSO4, 0.25mM of dNTPs, 0.5 µM of primers and 1U of Platinum Taq High Fidelity DNA Polymerase (Invitrogen 11304102) in a final volume of 40ul. The reaction included an initial denaturation step of 94 °C for 2min, followed by 45 cycles each including 94 °C for 20min, 55°C for 30min and 72 °C for 3min. After amplification, the PCR product was purified using the PCR Clean-up kit (Macherey-Nagel 740609.250) and quantified by Qubit dsDNA BR (Invitrogen Q33265). A total of 1µg of the purified PCR product was used for in vitro transcription using the HiScribe T7 High Yield RNA synthesis (NEB E2040S), as recommended by the manufacturer. Finally, the synthetic RNA was checked by electrophoresis and was quantified by nanodrop. In addition, to give a precise estimate of copies/µl of the synthetic RNA, serial 10-fold dilutions were quantified by digital PCR. At the beginning of the extraction, ∼8,000 copies of synthetic RNA were added to each well of the 96-well plate. For each experiment, synthetic RNA was added in three wells, containing the healthy donor cell cushion but not the participant-derived cells, just before the cDNA synthesis step, hence representing the 100% recovery control of RNA. We used two criteria to exclude wells with poor recovery: 1) wells with a recovery <30% to the average of the synthetic RNA control and/or 2) wells with a recovery inferior to the average copies of IC RNA across the entire plate + 2 standard deviation.

### Multiplex ELISA

The supernatants from re-stimulation experiments were collected, stored at -80°C, and analyzed by Legendplex Custom Panel (BioLegend) per the manufacturer’s protocol. Samples were acquired on an Intellicyt iQue Screener Plus (Sartorius). Data was analyzed by FlowJo v10.8.1 (BD Bioscience).

### RNA-sequencing

Bulk RNA sequencing was performed on samples from 6 study participants (P1, P4, P5, P6, P8, and P10). We sorted 10,000 live CD4^+^ T cells left untreated (NoTx), treated with anti-CD3/CD28, PMA/I, or autologous dendritic cells without addition of antigens (NoAg) or pulsed with cognate antigens. Due to technical challenges, the anti-CD3/CD28 sample from P1 and the NoAg sample from P6 were not collected. Cells were sorted in lysis buffer, spun and stored at -80°C until processed. RNA isolation was performed as previously described^11^. RNA quality was assessed using a Bioanalyzer RNA 6000 Nano Chip (Agilent Technologies). All samples had an RNA quality number (RQN) higher than 8, with a mean RIN of 9.89 (range 8.9-10). TrueSeq stranded mRNA libraries (Illumina) were prepared by the Single Cell Genomics Core at Johns Hopkins University and sequenced on an Illumina NovaSeq platform (1x100bp reads). Sequencing reads were aligned to the reference human genome using STAR (v2.7.10) and annotated using the hg19 build of the genome (GRCh37). Next, the number of reads mapping to each gene was quantified using RSEM (v1.3.3). RNA counts were imported into R (v4.1) where normalization for library size and regularized-logarithmic transformation of counts was performed using DESeq2 (v1.34). Differential expression analysis was performed with DESeq2. Differentially expressed genes were defined as any genes with absolute log-fold changes (LFC) larger than 1.2 at a false discovery rate (FDR) of 0.05. The list of genes targeted by the transcription factors (TF) NF-kB and NFAT was compiled based on previously published data^100–102^. Genes were selected if they are expressed in CD4^+^ T cells, contain a TF-binding site in their promoter and have been shown to directly interact with the TF (e.g., CHIP-seq data) ( https://www.bu.edu/nf-kb/gene-resources/target-genes/). Additional analysis and data visualization were performed with iDEP v1.1 (https://bmcbioinformatics.biomedcentral.com/articles/10.1186/s12859-018-2486-6).

#### TCRβ sequencing

gDNA was isolated from CD4^+^ T cells, quantified as described above, and diluted in Tris-acetate-EDTA to a concentration of 10 ng/μL (for a total of up to 1 μg per sample). TCRβ sequencing data were generated using the ImmunoSEQ hsTCRB assay, version 4, in survey mode (Adaptive Biotechnologies). Analyses on TCR data were conducted on the ImmuneAnalyzer platform (Adaptive Biotechnologies).

#### Quantification and statistical analyses

Descriptive statistics, tests for normality, 2-tailed Student’s t-test and one-way ANOVA tests were used to determine statistical significance using GraphPad Prism v9.0. P values lower than 0.05 were considered significant, unless otherwise stated.

#### Data availability

HIV-1 sequences are available on GenBank (sequences were submitted, accession numbers are pending). TCRβ sequencing data can be accessed through the ImmuneAccess database (doi pending). RNAseq data has been deposited to NCBI SRA and will be available after publication.

#### Study approval

The Johns Hopkins Institutional Review Board, the Wistar Institute, and the UCSF Committee on Human Research approved this study. All study participants provided written informed consent before enrollment.

## Notes

### Competing Interest Statement

Aspects of the intact proviral DNA assay (IPDA) are the subject of a patent application ('Compositions and methods Related to characterizing proviral reservoirs', PCT/US16/28822) filed by Johns Hopkins University, and RFS is one of the inventors on this application. The other authors have declared that no conflict of interest exists.

## References

1. Chun, T.W., Finzi, D., Margolick, J., Chadwick, K., Schwartz, D., and Siliciano, R.F. (1995). In vivo fate of HIV-1-infected T cells: Quantitative analysis of the transition to stable latency. Nat. Med. 1, 1284–1290. 10.1038/nm1295-1284.

2. Finzi, D., Hermankova, M., Pierson, T., Carruth, L.M., Buck, C., Chaisson, R.E., Quinn, T.C., Chadwick, K., Margolick, J., Brookmeyer, R., et al. (1997). Identification of a reservoir for HIV-1 in patients on highly active antiretroviral therapy. Science 278, 1295–1300. 10.1126/science.278.5341.1295.

3. Wong, J.K., Hezareh, M., Günthard, H.F., Havlir, D. V., Ignacio, C.C., Spina, C.A., and Richman, D.D. (1997). Recovery of Replication-Competent HIV Despite Prolonged Suppression of Plasma Viremia. Science 278, 1291–1295. 10.1126/SCIENCE.278.5341.1291.

4. Chun, T.W., Carruth, L., Finzi, D., Shen, X., DiGiuseppe, J.A., Taylor, H., Hermankova, M., Chadwick, K., Margolick, J., Quinn, T.C., et al. (1997). Quantification of latent tissue reservoirs and total body viral load in HIV-1 infection. Nature 387, 183–188. 10.1038/387183a0.

5. Berkhout, B., Silverman, R.H., and Jeang, K.T. (1989). Tat trans-activates the human immunodeficiency virus through a nascent RNA target. Cell 59, 273–282. 10.1016/0092-8674(89)90289-4.

6. Nabel, G., and Baltimore, D. (1987). An inducible transcription factor activates expression of human immunodeficiency virus in T cells. Nature 326, 711–713. 10.1038/326711a0.

7. Siliciano, R.F., and Greene, W.C. (2011). HIV Latency. Cold Spring Harb Perspect. Med. 1, a007096. 10.1101/CSHPERSPECT.A007096.

8. Chan, J.K.L., and Greene, W.C. (2011). NF-κB/Rel: Agonist and antagonist roles in HIV-1 latency. Curr. Opin. HIV AIDS 6, 12–18. 10.1097/COH.0b013e32834124fd.

9. Siekevitz, M., Josephs, S.F., Dukovich, M., Peffer, N., Wong-Staal, F., and Greene, W.C. (1987). Activation of the HIV-1 LTR by T cell mitogens and the trans-activator protein of HTLV-I. Science 238, 1575–1578. 10.1126/SCIENCE.2825351.

10. Crooks, A.M., Bateson, R., Cope, A.B., Dahl, N.P., Griggs, M.K., Kuruc, J.A.D., Gay, C.L., Eron, J.J., Margolis, D.M., Bosch, R.J., et al. (2015). Precise Quantitation of the Latent HIV-1 Reservoir: Implications for Eradication Strategies. J. Infect. Dis. 212, 1361–1365. 10.1093/INFDIS/JIV218.

11. Siliciano, J.D., Kajdas, J., Finzi, D., Quinn, T.C., Chadwick, K., Margolick, J.B., Kovacs, C., Gange, S.J., and Siliciano, R.F. (2003). Long-term follow-up studies confirm the stability of the latent reservoir for HIV-1 in resting CD4+ T cells. Nat. Med. 9, 727–728. 10.1038/nm880.

12. McMyn, N.F., Varriale, J., Fray, E.J., Zitzmann, C., MacLeod, H., Lai, J., Singhal, A., Moskovljevic, M., Garcia, M.A., Lopez, B.M., et al. (2023). The latent reservoir of inducible, infectious HIV-1 does not decrease despite decades of antiretroviral therapy. J. Clin. Invest. 133. 10.1172/JCI171554.

13. Davey, R.T., Bhat, N., Yoder, C., Chun, T.W., Metcalf, J.A., Dewar, R., Natarajan, V., Lempicki, R.A., Adelsberger, J.W., Miller, K.D., et al. (1999). HIV-1 and T cell dynamics after interruption of highly active antiretroviral therapy (HAART) in patients with a history of sustained viral suppression. Proc. Natl. Acad. Sci. USA 96, 15109–15114. 10.1073/PNAS.96.26.15109.

14. Rothenberger, M.K., Keele, B.F., Wietgrefe, S.W., Fletcher, C. V., Beilman, G.J., Chipman, J.G., Khoruts, A., Estes, J.D., Anderson, J., Callisto, S.P., et al. (2015). Large number of rebounding/founder HIV variants emerge from multifocal infection in lymphatic tissues after treatment interruption. Proc. Natl. Acad. Sci. USA 112, E1126–E1134.

15. Bui, J.K., Sobolewski, M.D., Keele, B.F., Spindler, J., Musick, A., Wiegand, A., Luke, B.T., Shao, W., Hughes, S.H., Coffin, J.M., et al. (2017). Proviruses with identical sequences comprise a large fraction of the replication-competent HIV reservoir. PLoS Pathog. 13, e1006283. 10.1371/JOURNAL.PPAT.1006283.

16. Hosmane, N.N., Kwon, K.J., Bruner, K.M., Capoferri, A.A., Beg, S., Rosenbloom, D.I.S., Keele, B.F., Ho, Y.C., Siliciano, J.D., and Siliciano, R.F. (2017). Proliferation of latently infected CD4 + T cells carrying replication-competent HIV-1: Potential role in latent reservoir dynamics. J. Exp. Med. 214, 959–972.

17. Bailey, J.R., Sedaghat, A.R., Kieffer, T., Brennan, T., Lee, P.K., Wind-Rotolo, M., Haggerty, C.M., Kamireddi, A.R., Liu, Y., Lee, J., et al. (2006). Residual Human Immunodeficiency Virus Type 1 Viremia in Some Patients on Antiretroviral Therapy Is Dominated by a Small Number of Invariant Clones Rarely Found in Circulating CD4+ T Cells. J. Virol. 80, 6441–6457. 10.1128/jvi.00591-06.

18. Wagner, T.A., McLaughlin, S., Garg, K., Cheung, C.Y.K., Larsen, B.B., Styrchak, S., Huang, H.C., Edlefsen, P.T., Mullins, J.I., and Frenkel, L.M. (2014). Proliferation of cells with HIV integrated into cancer genes contributes to persistent infection. Science 345, 570–573.

19. Maldarelli, F., Wu, X., Su, L., Simonetti, F.R., Shao, W., Hill, S., Spindler, J., Ferris, A.L., Mellors, J.W., Kearney, M.F., et al. (2014). Specific HIV integration sites are linked to clonal expansion and persistence of infected cells. Science 345, 179–183.

20. Simonetti, F.R., Zhang, H., Soroosh, G.P., Duan, J., Rhodehouse, K., Hill, A.L., Beg, S.A., McClurkan, K., Raymond, H.E., Nobles, C.L., et al. (2021). Antigen-driven clonal selection shapes the persistence of HIV-1–infected CD4+ T cells in vivo. J. Clin. Invest. 131. 10.1172/JCI145254.

21. Simonetti, F.R., Sobolewski, M.D., Fyne, E., Shao, W., Spindler, J., Hattori, J., Anderson, E.M., Watters, S.A., Hill, S., Wu, X., et al. (2016). Clonally expanded CD4+ T cells can produce infectious HIV-1 in vivo. Proc. Natl. Acad. Sci. USA 113, 1883–1888. 10.1073/PNAS.1522675113.

22. Cohn, L.B., Da Silva, I.T., Valieris, R., Huang, A.S., Lorenzi, J.C.C., Cohen, Y.Z., Pai, J.A., Butler, A.L., Caskey, M., Jankovic, M., et al. (2018). Clonal CD4+ T cells in the HIV-1 latent reservoir display a distinct gene profile upon reactivation. Nat. Med. 24, 604. 10.1038/S41591-018-0017-7.

23. Tobin, N.H., Learn, G.H., Holte, S.E., Wang, Y., Melvin, A.J., McKernan, J.L., Pawluk, D.M., Mohan, K.M., Lewis, P.F., Mullins, J.I., et al. (2005). Evidence that low-level viremias during effective highly active antiretroviral therapy result from two processes: expression of archival virus and replication of virus. J. Virol. 79, 9625–9634. 10.1128/JVI.79.15.9625-9634.2005.

24. Mendoza, P., Jackson, J.R., Oliveira, T.Y., Gaebler, C., Ramos, V., Caskey, M., Jankovic, M., Nussenzweig, M.C., and Cohn, L.B. (2020). Antigen-responsive cd4+ t cell clones contribute to the hiv-1 latent reservoir. J. Exp. Med. 217. 10.1084/JEM.20200051/151689.

25. Collora, J.A., Liu, R., Pinto-Santini, D., Ravindra, N., Ganoza, C., Lama, J.R., Alfaro, R., Chiarella, J., Spudich, S., Mounzer, K., et al. (2022). Single-cell multiomics reveals persistence of HIV-1 in expanded cytotoxic T cell clones. Immunity 55, 1013–1031.e7. 10.1016/J.IMMUNI.2022.03.004.

26. Chomont, N., El-Far, M., Ancuta, P., Trautmann, L., Procopio, F.A., Yassine-Diab, B., Boucher, G., Boulassel, M.R., Ghattas, G., Brenchley, J.M., et al. (2009). HIV reservoir size and persistence are driven by T cell survival and homeostatic proliferation. Nat. Med. 15, 893–900. 10.1038/NM.1972.

27. Coffin, J.M., Bale, M.J., Wells, D., Guo, S., Luke, B., Zerbato, J.M., Sobolewski, M.D., Sia, T., Shao, W., Wu, X., et al. (2021). Integration in oncogenes plays only a minor role in determining the in vivo distribution of HIV integration sites before or during suppressive antiretroviral therapy. PLoS Pathog. 17, e1009141. 10.1371/JOURNAL.PPAT.1009141.

28. Henrich, T.J., Hobbs, K.S., Hanhauser, E., Scully, E., Hogan, L.E., Robles, Y.P., Leadabrand, K.S., Marty, F.M., Palmer, C.D., Jost, S., et al. (2017). Human Immunodeficiency Virus Type 1 Persistence Following Systemic Chemotherapy for Malignancy. J. Infect. Dis. 216, 254–262.

29. Wei, X., Ghosh, S.K., Taylor, M.E., Johnson, V.A., Emini, E.A., Deutsch, P., Lifson, J.D., Bonhoeffer, S., Nowak, M.A., Hahn, B.H., et al. (1995). Viral dynamics in human immunodeficiency virus type 1 infection. Nature 1995 373:6510 *373*, 117–122. 10.1038/373117a0.

30. Koup, R.A., Safrit, J.T., Cao, Y., Andrews, C.A., Mcleod, G., Borkowsky, W., Farthing, C., and Ho, D.D. (1994). Temporal Association of Cellular Immune Responses with the Initial Control of Viremia in Primary Human Immunodeficiency Virus Type 1 Syndrome. J. Virol. 68, 4650–4655.

31. Ahmed, R., and Gray, D. (1996). Immunological Memory and Protective Immunity: Understanding Their Relation. Science 272, 54–60. 10.1126/SCIENCE.272.5258.54.

32. Kwon, K.J., Timmons, A.E., Sengupta, S., Simonetti, F.R., Zhang, H., Hoh, R., Deeks, S.G., Siliciano, J.D., and Siliciano, R.F. (2020). Different human resting memory CD4+ T cell subsets show similar low inducibility of latent HIV-1 proviruses. Sci. Transl. Med. 12 (528): eaax6795.

33. Bruner, K.M., Murray, A.J., Pollack, R.A., Soliman, M.G., Laskey, S.B., Capoferri, A.A., Lai, J., Strain, M.C., Lada, S.M., Hoh, R., et al. (2016). Defective proviruses rapidly accumulate during acute HIV-1 infection. Nat. Med. 2016 22:9 22, 1043–1049. 10.1038/nm.4156.

34. Eriksson, S., Graf, E.H., Dahl, V., Strain, M.C., Yukl, S.A., Lysenko, E.S., Bosch, R.J., Lai, J., Chioma, S., Emad, F., et al. (2013). Comparative Analysis of Measures of Viral Reservoirs in HIV-1 Eradication Studies. PLoS Pathog. 9, e1003174. 10.1371/JOURNAL.PPAT.1003174.

35. Betts, M.R., Ambrozak, D.R., Douek, D.C., Bonhoeffer, S., Brenchley, J.M., Casazza, J.P., Koup, R.A., and Picker, L.J. (2001). Analysis of total human immunodeficiency virus (HIV)-specific CD4(+) and CD8(+) T-cell responses: relationship to viral load in untreated HIV infection. J. Virol. 75, 11983– 11991. 10.1128/JVI.75.24.11983-11991.2001.

36. Bacher, P., Schink, C., Teutschbein, J., Kniemeyer, O., Assenmacher, M., Brakhage, A.A., and Scheffold, A. (2013). Antigen-reactive T cell enrichment for direct, high-resolution analysis of the human naive and memory Th cell repertoire. J. Immunol. 190, 3967–3976.

37. Cillo, A.R., Sobolewski, M.D., Bosch, R.J., Fyne, E., Piatak, M., Coffin, J.M., and Mellors, J.W. (2014). Quantification of HIV-1 latency reversal in resting CD4+ T cells from patients on suppressive antiretroviral therapy. Proc. Natl. Acad. Sci. USA 111, 7078–7083.

38. Kim, Y., Anderson, J.L., and Lewin, S.R. (2018). Getting the “Kill” into “Shock and Kill”: Strategies to Eliminate Latent HIV. Cell Host Microbe 23, 14–26. 10.1016/j.chom.2017.12.004.

39. Feng, Y., Reinherz, E.L., and Lang, M.J. (2018). αβ T Cell Receptor Mechanosensing Forces out Serial Engagement. Trends Immunol. 39, 596–609. 10.1016/J.IT.2018.05.005.

40. Plantin, J., Massanella, M., and Chomont, N. (2018). Inducible HIV RNA transcription assays to measure HIV persistence: Pros and cons of a compromise. Retrovirology 15, 1–11.

41. Courtney, A.H., Lo, W.L., and Weiss, A. (2018). TCR Signaling: Mechanisms of Initiation and Propagation. Trends Biochem. Sci. 43, 108–123. 10.1016/J.TIBS.2017.11.008.

42. Vollbrecht, T., Angerstein, A.O., Menke, B., Kumar, N.M., de Oliveira, M.F., Richman, D.D., and Guatelli, J.C. (2020). Inconsistent reversal of HIV-1 latency ex vivo by antigens of HIV-1, CMV, and other infectious agents. Retrovirology 17, 1–12.

43. Kristoff, J., Palma, M.L., Garcia-Bates, T.M., Shen, C., Sluis-Cremer, N., Gupta, P., Rinaldo, C.R., and Mailliard, R.B. (2019). Type 1-programmed dendritic cells drive antigen-specific latency reversal and immune elimination of persistent HIV-1. EBioMedicine 43, 295–306. 10.1016/j.ebiom.2019.03.077.

44. Bacher, P., Hohnstein, T., Beerbaum, E., Röcker, M., Blango, M.G., Kaufmann, S., Röhmel, J., Eschenhagen, P., Grehn, C., Seidel, K., et al. (2019). Human Anti-fungal Th17 Immunity and Pathology Rely on Cross-Reactivity against Candida albicans. Cell 176, 1340–1355.e15. 10.1016/j.cell.2019.01.041.

45. Frentsch, M., Arbach, O., Kirchhoff, D., Moewes, B., Worm, M., Rothe, M., Scheffold, A., and Thiel, A. (2005). Direct access to CD4+ T cells specific for defined antigens according to CD154 expression. Nat. Med. 2005 11:10 *11*, 1118–1124. 10.1038/nm1292.

46. Saggau, C., Martini, G.R., Rosati, E., Meise, S., Messner, B., Kamps, A.K., Bekel, N., Gigla, J., Rose, R., Voß, M., et al. (2022). The pre-exposure SARS-CoV-2-specific T cell repertoire determines the quality of the immune response to vaccination. Immunity 55, 1924–1939.e5. 10.1016/J.IMMUNI.2022.08.003.

47. Martini, G.R., Tikhonova, E., Rosati, E., DeCelie, M.B., Sievers, L.K., Tran, F., Lessing, M., Bergfeld, A., Hinz, S., Nikolaus, S., et al. (2023). Selection of cross-reactive T cells by commensal and food-derived yeasts drives cytotoxic TH1 cell responses in Crohn’s disease. Nat. Med. 10.1038/S41591-023-02556-5.

48. Morisita M (1959). Measuring of the Dispersion of Individuals and Analysis of the Distributional Patterns. - Memories of Faculty Science, Kyushu University, Series E. Biology 2: 215–235.

49. Bruner, K.M., Wang, Z., Simonetti, F.R., Bender, A.M., Kwon, K.J., Sengupta, S., Fray, E.J., Beg, S.A., Antar, A.A.R., Jenike, K.M., et al. (2019). A quantitative approach for measuring the reservoir of latent HIV-1 proviruses. Nature 566, 120–125. 10.1038/s41586-019-0898-8.

50. Simonetti, F.R., White, J.A., Tumiotto, C., Ritter, K.D., Cai, M., Gandhi, R.T., Deeks, S.G., Howell, B.J., Montaner, L.J., Blankson, J.N., et al. (2020). Intact proviral DNA assay analysis of large cohorts of people with HIV provides a benchmark for the frequency and composition of persistent proviral DNA. Proc. Natl. Acad. Sci. USA 117, 18692–18700. 10.1073/PNAS.2006816117.

51. Kufera, J.T., Armstrong, C., Wu, F., Singhal, A., Zhang, H., Lai, J., Wilkins, H.N., Simonetti, F.R., Siliciano, J.D., and Siliciano, R.F. (2024). CD4+ T cells with latent HIV-1 have reduced proliferative responses to T cell receptor stimulation. J. Exp. Med. 221. 10.1084/JEM.20231511/276522.

52. McDonald, D., Wu, L., Bohks, S.M., KewalRamani, V.N., Unutmaz, D., and Hope, T.J. (2003). Recruitment of HIV and its receptors to dendritic cell-T cell junctions. Science 300, 1295–1297. 10.1126/science.1084238.

53. Archin, N.M., and Margolis, D.M. (2014). Emerging strategies to deplete the HIV reservoir. Curr. Opin. Infect. Dis. 27: 29–35, 29. 10.1097/QCO.0000000000000026.

54. Karn, J., and Stoltzfus, C.M. (2012). Transcriptional and posttranscriptional regulation of HIV-1 gene expression. Cold Spring Harb. Perspect. Med. 2, a006916. 10.1101/cshperspect.a006916.

55. Hiscott, J., Kwon, H., and Génin, P. (2001). Hostile takeovers: viral appropriation of the NF-kappaB pathway. J. Clin. Invest. 107, 143–151. 10.1172/JCI11918.

56. Van Lint, C., Bouchat, S., and Marcello, A. (2013). HIV-1 transcription and latency: An update. Retrovirology 10, 1–38. 10.1186/1742-4690-10-67.

57. Morou, A., Brunet-Ratnasingham, E., Dubé, M., Charlebois, R., Mercier, E., Darko, S., Brassard, N., Nganou-Makamdop, K., Arumugam, S., Gendron-Lepage, G., et al. (2019). Altered differentiation is central to HIV-specific CD4+ T cell dysfunction in progressive disease. Nat. Immunol. 20, 1059– 1070. 10.1038/S41590-019-0418-X.

58. Reed, J.C. (1999). Bcl-2 family proteins. Oncogene 1998 17:25 17, 3225–3236. 10.1038/sj.onc.1202591.

59. Gavet, O., and Pines, J. (2010). Progressive Activation of CyclinB1-Cdk1 Coordinates Entry to Mitosis. Dev. Cell. 18, 533–543. 10.1016/J.DEVCEL.2010.02.013.

60. Yukl, S.A., Kaiser, P., Kim, P., Telwatte, S., Joshi, S.K., Vu, M., Lampiris, H., and Wong, J.K. (2018). HIV latency in isolated patient CD4+ T cells may be due to blocks in HIV transcriptional elongation, completion, and splicing. Sci. Transl. Med. 10. (430):eaap9927.

61. Telwatte, S., Morón-López, S., Aran, D., Kim, P., Hsieh, C., Joshi, S., Montano, M., Greene, W.C., Butte, A.J., Wong, J.K., et al. (2019). Heterogeneity in HIV and cellular transcription profiles in cell line models of latent and productive infection: implications for HIV latency. Retrovirology 16. 10.1186/S12977-019-0494-X.

62. Shan, L., Rabi, S.A., Laird, G.M., Eisele, E.E., Zhang, H., Margolick, J.B., and Siliciano, R.F. (2013). A Novel PCR Assay for Quantification of HIV-1 RNA. J. Virol. 87, 6521–6525. 10.1128/JVI.00006-13/FORMAT/EPUB.

63. Bullen, C.K., Laird, G.M., Durand, C.M., Siliciano, J.D., and Siliciano, R.F. (2014). New ex vivo approaches distinguish effective and ineffective single agents for reversing HIV-1 latency in vivo. Nat. Med. 20, 425–429. 10.1038/nm.3489.

64. Laird, G.M., Bullen, C.K., Rosenbloom, D.I.S., Martin, A.R., Hill, A.L., Durand, C.M., Siliciano, J.D., and Siliciano, R.F. (2015). Ex vivo analysis identifies effective HIV-1 latency-reversing drug combinations. J. Clin. Invest. 125, 1901–1912. 10.1172/JCI80142.

65. Rosalia, R.A., Quakkelaar, E.D., Redeker, A., Khan, S., Camps, M., Drijfhout, J.W., Silva, A.L., Jiskoot, W., van Hall, T., van Veelen, P.A., et al. (2013). Dendritic cells process synthetic long peptides better than whole protein, improving antigen presentation and T-cell activation. Eur. J. Immunol. 43, 2554–2565. 10.1002/EJI.201343324.

66. Dragoni, F., Kwaa, A.K., Traut, C.C., Veenhuis, R.T., Woldemeskel, B.A., Camilo-Contreras, A., Raymond, H.E., Dykema, A.G., Scully, E.P., Rosecrans, A.M., et al. (2023). Proviral location affects cognate peptide–induced virus production and immune recognition of HIV-1–infected T cell clones. J. Clin. Invest. 133. 10.1172/JCI171097.

67. Pollack, R.A., Jones, R.B., Pertea, M., Bruner, K.M., Martin, A.R., Thomas, A.S., Capoferri, A.A., Beg, S.A., Huang, S.H., Karandish, S., et al. (2017). Defective HIV-1 Proviruses Are Expressed and Can Be Recognized by Cytotoxic T Lymphocytes, which Shape the Proviral Landscape. Cell Host Microbe 21, 494–506.e4. 10.1016/J.CHOM.2017.03.008.

68. Scrimieri, F., Bastian, E., Smith, M., Rehm, C.A., Morse, C., Kuruppu, J., McLaughlin, M., Chang, W., Sereti, I., Kovacs, J.A., et al. (2024). Transcriptionally Active Defective HIV-1 Proviruses and Their Association with Immunological Nonresponse to Antiretroviral Therapy. J. Infect. Dis. 10.1093/INFDIS/JIAE009.

69. Dubé, M., Tastet, O., Dufour, C., Sannier, G., Brassard, N., Delgado, G.G., Pagliuzza, A., Richard, C., Nayrac, M., Routy, J.P., et al. (2023). Spontaneous HIV expression during suppressive ART is associated with the magnitude and function of HIV-specific CD4+ and CD8+ T cells. Cell Host Microbe 31, 1507–1522.e5. 10.1016/J.CHOM.2023.08.006.

70. Takata, H., Mitchell, J.L., Pacheco, J., Pagliuzza, A., Pinyakorn, S., Buranapraditkun, S., Sacdalan, C., Leyre, L., Nathanson, S., Kakazu, J.C., et al. (2023). An active HIV reservoir during ART is associated with maintenance of HIV-specific CD8+ T cell magnitude and short-lived differentiation status. Cell Host Microbe 31, 1494–1506.e4. 10.1016/J.CHOM.2023.08.012.

71. Turpin, J., Yurick, D., Khoury, G., Pham, H., Locarnini, S., Melamed, A., Witkover, A., Wilson, K., Purcell, D., Bangham, C.R.M., et al. (2019). Impact of Hepatitis B Virus Coinfection on Human T-Lymphotropic Virus Type 1 Clonality in an Indigenous Population of Central Australia. J. Infect. Dis. 219, 562–567. 10.1093/INFDIS/JIY546.

72. Elliott, J.H., Wightman, F., Solomon, A., Ghneim, K., Ahlers, J., Cameron, M.J., Smith, M.Z., Spelman, T., McMahon, J., Velayudham, P., et al. (2014). Activation of HIV Transcription with Short-Course Vorinostat in HIV-Infected Patients on Suppressive Antiretroviral Therapy. PLoS Pathog. 10, e1004473. 10.1371/JOURNAL.PPAT.1004473.

73. Janjic, A., Wange, L.E., Bagnoli, J.W., Geuder, J., Nguyen, P., Richter, D., Vieth, B., Vick, B., Jeremias, I., Ziegenhain, C., et al. (2022). Prime-seq, efficient and powerful bulk RNA sequencing. Genome Biol. 23. 10.1186/S13059-022-02660-8.

74. Einkauf, K.B., Osborn, M.R., Gao, C., Sun, W., Sun, X., Lian, X., Parsons, E.M., Gladkov, G.T., Seiger, K.W., Blackmer, J.E., et al. (2022). Parallel analysis of transcription, integration, and sequence of single HIV-1 proviruses. Cell 185, 266–282.e15. 10.1016/J.CELL.2021.12.011.

75. Jordan, A., Bisgrove, D., and Verdin, E. (2003). HIV reproducibly establishes a latent infection after acute infection of T cells in vitro. EMBO J. 22, 1868–1877. 10.1093/EMBOJ/CDG188.

76. Folks, T.M., Clouse, K.A., Justement, J., Rabson, A., Duh, E., Kehrl, J.H., and Fauci, A.S. (1989). Tumor necrosis factor α induces expression of human immunodeficiency virus in a chronically infected T-cell clone. Proc. Natl. Acad. Sci. USA 86, 2365–2368. 10.1073/PNAS.86.7.2365.

77. Siliciano, J.D., and Siliciano, R.F. (2021). Low Inducibility of Latent Human Immunodeficiency Virus Type 1 Proviruses as a Major Barrier to Cure. J. Infect. Dis. 223, S13–S21. 10.1093/INFDIS/JIAA649.

78. Liu, R., Simonetti, F.R., and Ho, Y.C. (2020). The forces driving clonal expansion of the HIV-1 latent reservoir. Virol. J. 2020 17:1 *17*, 1–13. 10.1186/S12985-019-1276-8.

79. Wang, Z., Gurule, E.E., Brennan, T.P., Gerold, J.M., Kwon, K.J., Hosmane, N.N., Kumar, M.R., Beg, S.A., Capoferri, A.A., Ray, S.C., et al. (2018). Expanded cellular clones carrying replication-competent HIV-1 persist, wax, and wane. Proc. Natl. Acad. Sci. USA 115, E2575–E2584. 10.1073/PNAS.1720665115.

80. Grakoui, A., Bromley, S.K., Sumen, C., Davis, M.M., Shaw, A.S., Allen, P.M., and Dustin, M.L. (1999). The immunological synapse: A molecular machine controlling T cell activation. Science 285, 221–227. 10.1126/SCIENCE.285.5425.221.

81. Board, N.L., Moskovljevic, M., Wu, F., Siliciano, R.F., and Siliciano, J.D. (2021). Engaging innate immunity in HIV-1 cure strategies. Nat. Rev. Immunol. 22, 499–512. 10.1038/s41577-021-00649-1.

82. Casazza, J.P., Brenchley, J.M., Hill, B.J., Ayana, R., Ambrozak, D., Roederer, M., Douek, D.C., Betts, M.R., and Koup, R.A. (2009). Autocrine Production of β-Chemokines Protects CMV-Specific CD4+ T Cells from HIV Infection. PLoS Pathog. 5, e1000646. 10.1371/JOURNAL.PPAT.1000646.

83. Douek, D.C., Brenchley, J.M., Betts, M.R., Ambrozak, D.R., Hill, B.J., Okamoto, Y., Casazza, J.P., Kuruppu, J., Kunstman, K., Wolinsky, S., et al. (2002). HIV preferentially infects HIV-specific CD4+ T cells. Nature 417, 95–98. 10.1038/417095A.

84. Gantner, P., Buranapraditkun, S., Pagliuzza, A., Dufour, C., Pardons, M., Mitchell, J.L., Kroon, E., Sacdalan, C., Tulmethakaan, N., Pinyakorn, S., et al. (2023). HIV rapidly targets a diverse pool of CD4+ T cells to establish productive and latent infections. Immunity 56, 653–668.e5. 10.1016/J.IMMUNI.2023.01.030.

85. Shaw, J.P., Utz, P.J., Durand, D.B., Toole, J.J., Emmel, E.A., and Crabtree, G.R. (1988). Identification of a putative regulator of early T cell activation genes. Science 241, 4972–4975. 10.1126/SCIENCE.3260404.

86. Pahl, H.L. (1999). Activators and target genes of Rel/NF-κB transcription factors. Oncogene 1999 18:49 18, 6853–6866. 10.1038/sj.onc.1203239.

87. Brignall, R., Cauchy, P., Bevington, S.L., Gorman, B., Pisco, A.O., Bagnall, J., Boddington, C., Rowe, W., England, H., Rich, K., et al. (2017). Integration of Kinase and Calcium Signaling at the Level of Chromatin Underlies Inducible Gene Activation in T Cells. J. Immunol. 199, 2652–2667. 10.4049/JIMMUNOL.1602033.

88. Archin, N.M., Liberty, A.L., Kashuba, A.D., Choudhary, S.K., Kuruc, J.D., Crooks, A.M., Parker, D.C., Anderson, E.M., Kearney, M.F., Strain, M.C., et al. (2012). Administration of vorinostat disrupts HIV-1 latency in patients on antiretroviral therapy. Nature 487, 482–485. 10.1038/nature11286.

89. Kjær, K., Leth, S., Konrad, C. V., Gunst, J.D., Nymann, R., Østergaard, L., Søgaard, O.S., Schleimann, M.H., Tolstrup, M., and Denton, P.W. (2021). Modest de novo Reactivation of Single HIV-1 Proviruses in Peripheral CD4+ T Cells by Romidepsin. Front. Virol. 1, 736395. 10.3389/FVIRO.2021.736395.

90. Pardons, M., Cole, B., Lambrechts, L., van Snippenberg, W., Rutsaert, S., Noppe, Y., De Langhe, N., Dhondt, A., Vega, J., Eyassu, F., et al. (2023). Potent latency reversal by Tat RNA-containing nanoparticle enables multi-omic analysis of the HIV-1 reservoir. Nat. Commun. 14. 10.1038/S41467-023-44020-5.

91. Lian, X., Seiger, K.W., Parsons, E.M., Gao, C., Sun, W., Gladkov, G.T., Roseto, I.C., Einkauf, K.B., Osborn, M.R., Chevalier, J.M., et al. (2023). Progressive transformation of the HIV-1 reservoir cell profile over two decades of antiviral therapy. Cell Host Microbe 31, 83–96.e5. 10.1016/J.CHOM.2022.12.002.

92. Huang, A.S., Ramos, V., Oliveira, T.Y., Gaebler, C., Jankovic, M., Nussenzweig, M.C., and Cohn, L.B. (2021). Integration features of intact latent HIV-1 in CD4+ T cell clones contribute to viral persistence. J. Exp. Med. 218. 10.1084/JEM.20211427.

93. Singh, K., Natarajan, V., Dewar, R., Rupert, A., Badralmaa, Y., Zhai, T., Winchester, N., Scrimieri, F., Smith, M., Davis, I., et al. (2023). Long-term persistence of transcriptionally active “defective” HIV-1 proviruses: implications for persistent immune activation during antiretroviral therapy. AIDS 37, 2119–2130. 10.1097/QAD.0000000000003667.

94. Wu, F., and Simonetti, F.R. (2023). Learning from Persistent Viremia: Mechanisms and Implications for Clinical Care and HIV-1 Cure. Curr HIV/AIDS Rep. 20, 428. 10.1007/S11904-023-00674-W.

95. White, J.A., Wu, F., Yasin, S., Moskovljevic, M., Varriale, J., Dragoni, F., Camilo-Contreras, A., Duan, J., Zheng, M.Y., Tadzong, N.F., et al. (2023). Clonally expanded HIV-1 proviruses with 5′-leader defects can give rise to nonsuppressible residual viremia. Journal of Clinical Investigation 133. 10.1172/JCI165245

96. Kaufmann, D.E., Bailey, P.M., Sidney, J., Wagner, B., Norris, P.J., Johnston, M.N., Cosimi, L.A., Addo, M.M., Lichterfeld, M., Altfeld, M., et al. (2004). Comprehensive Analysis of Human Immunodeficiency Virus Type 1-Specific CD4 Responses Reveals Marked Immunodominance of gag and nef and the Presence of Broadly Recognized Peptides. J. Virol. 78, 4463–4477. 10.1128/JVI.78.9.4463-4477.2004

97. Kearney, M.F., Spindler, J., Shao, W., Yu, S., Anderson, E.M., O’Shea, A., Rehm, C., Poethke, C., Kovacs, N., Mellors, J.W., et al. (2014). Lack of Detectable HIV-1 Molecular Evolution during Suppressive Antiretroviral Therapy. PLoS Pathog 10, e1004010. 10.1371/JOURNAL.PPAT.1004010.

98. Thompson, J.D., Higgins, D.G., and Gibson, T.J. (1994). CLUSTAL W: improving the sensitivity of progressive multiple sequence alignment through sequence weighting, position-specific gap penalties and weight matrix choice. Nucleic Acids Res 22, 4673–4680. 10.1093/NAR/22.22.4673.

99. Kumar, S., Stecher, G., and Tamura, K. (2016). MEGA7: Molecular Evolutionary Genetics Analysis Version 7.0 for Bigger Datasets. Mol Biol Evol 33, 1870–1874. 10.1093/MOLBEV/MSW054.

100. Liu, T., Zhang, L., Joo, D., and Sun, S.C. (2017). NF-κB signaling in inflammation. Signal Transduction and Targeted Therapy 2017 2:1 2, 1–9. 10.1038/sigtrans.2017.23.

101. Hermann-Kleiter, N., and Baier, G. (2010). NFAT pulls the strings during CD4+ T helper cell effector functions. Blood 115, 2989–2997. 10.1182/BLOOD-2009-10-233585.

102. Elliot, T.A.E., Jennings, E.K., Lecky, D.A.J., Thawait, N., Flores-Langarica, A., Copland, A., Maslowski, K.M., Wraith, D.C., and Bending, D. (2021). Antigen and checkpoint receptor engagement recalibrates T cell receptor signal strength. Immunity 54, 2481–2496.e6. 10.1016/j.immuni.2021.08.020.

