## Supplemental Materials for "Cognate Antigen Engagement Induces HIV-1 Expression In CD4^+^ T Cells From People On Long-Term ART"

### SUPPLEMENTARY MATERIALS

**Table of contents**

**Supplementary Figures**

**Supplementary Figure S1.** TCRβ abundance and infection frequency within expanded Ag-responding cells. Related to Figure 1.

**Supplementary Figure S2.** Characterization of memory and activation status of expanded Ag-responding cells and sorting strategy upon different types of stimulating conditions. Related to Figure 2.

**Supplemental Figure S3.** Additional analyses on gene and protein expression upon stimulation with cognate Ag versus PMA/I. Related to Figure 3.

**Supplementary Figure S4.** Additional analyses of HIV-1 RNA expression upon re-stimulation. Related to Figure 4.

**Supplementary Figure S5.** Phylogenetic analysis of cell-associated HIV-1 RNA sequences from participant P7. Related to Figure 5.

**Supplementary Figure S6.** Validation experiments and additional analyses related to the limiting-dilution viral quantification assay. Related to Figure 6.

**Supplementary Tables**

**Supplementary Table S1.** Participant characteristics.

**Supplementary Table S2**. Primers and probes used in this study.

**Supplementary Table S3**. Sequence of the internal control used for the limiting dilution HIV-1 RNA assay.

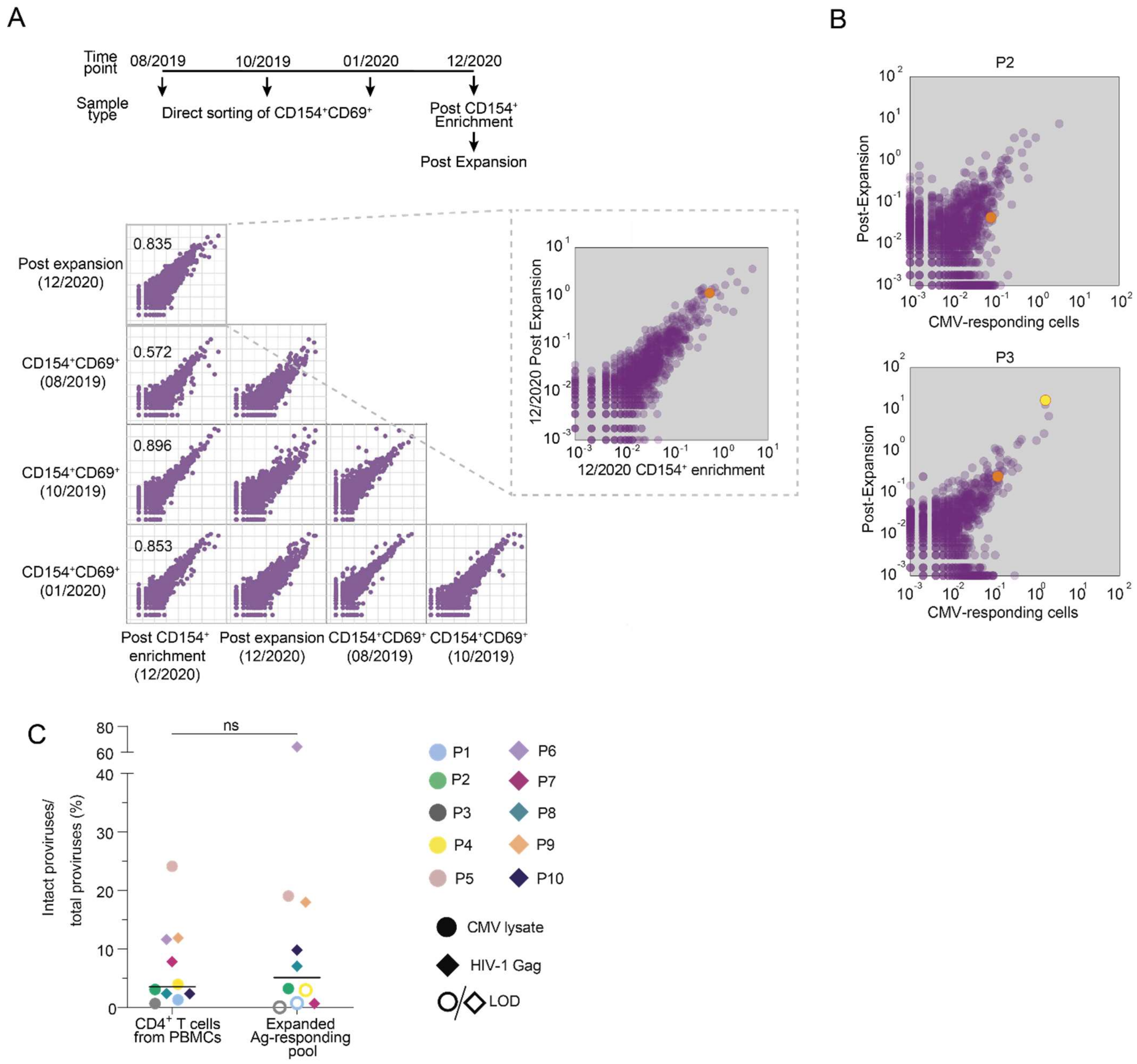


**Supplementary Figure S1**. **TCRβ abundance and infection frequency within expanded Ag-responding cells**. (**A**) Comparison of TCRβ frequencies among Ag-responsive cells obtained by magnetic enrichment (CD154), enrichment and expansion, or direct sorting of cells based on CD154 and CD69 expression from participant P1 at different time points. Numbers in the upper right corner of quadrants indicate the Morisita overlap index (see methods). Symbols indicate the percentage abundances of TCRβ sequences in two different samples, on a log10 scale. (**B**) Comparison of TCRβ sequence abundance among Ag-responding cells at the end of expansion and among cells sorted after overnight PBMC stimulation from Simonetti et al[6]; correlation plots from two representative participants are shown; colored circles indicate clonotypes carrying proviruses, previously characterized[6]. (**C**) Percentage of intact proviruses within purified $CD4^+$ T cells from PBMCs and expanded pools of Ag-responding cells. Horizontal bars represent the median value. Statistical significance was determined by paired t-test.

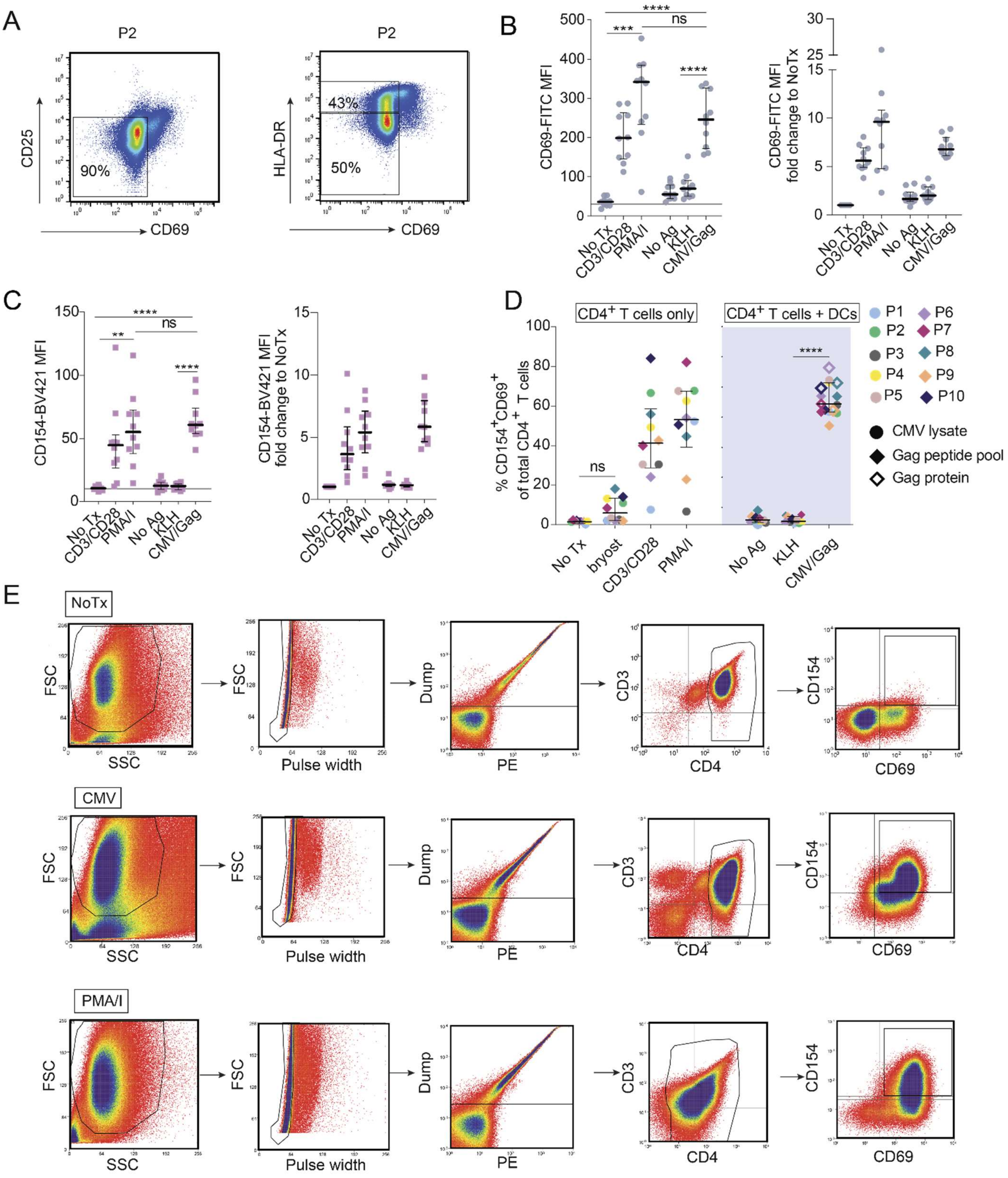


**Supplementary Figure S2. Characterization of memory and activation status of expanded Ag-responding cells and sorting strategy upon different types of stimulating conditions.** (**A**) Representative flow plots for CD25, HLA-DR, and CD69 expression on CD4⁺ T cells after expansion and 3 days of rest for participant P2. (**B**) Mean fluorescence intensity (MFI) of CD69 across all participants expressed as raw values (left graph). MFI of CD69 across all participants expressed as and as fold change to NoTx (right graph). (**C**) MFI of CD154 across all participants express as raw value (left graph). MFI of CD154 across all participants expressed as fold change to NoTx (right graph). (**D**) Percentage of C154$^{+}$CD69$^{+}$CD4$^{+}$ T cells in expanded pool of CMV-responding cells and Gag-responding cells upon different stimulating conditions. (**A-D**) All graphs are shown with horizontal bars representing median with interquartile range. Statistical significance (graphs B-D) was determined by using one-way ANOVA by Šídák's test for multiple comparisons, ** $P<0.01$, ***$P<0.001$, ****$P<0.0001$, ns- not statistically significant. (**E**) Representative flow plots showing gating sorting strategy for NoTx, CMV and PMA/I treatment conditions. NoTx- no treatment, CD3/CD28- anti-CD3/CD28 activating, PMA/I- pma/ionomycin, bryost- bryostatin, NoAg- no antigen.

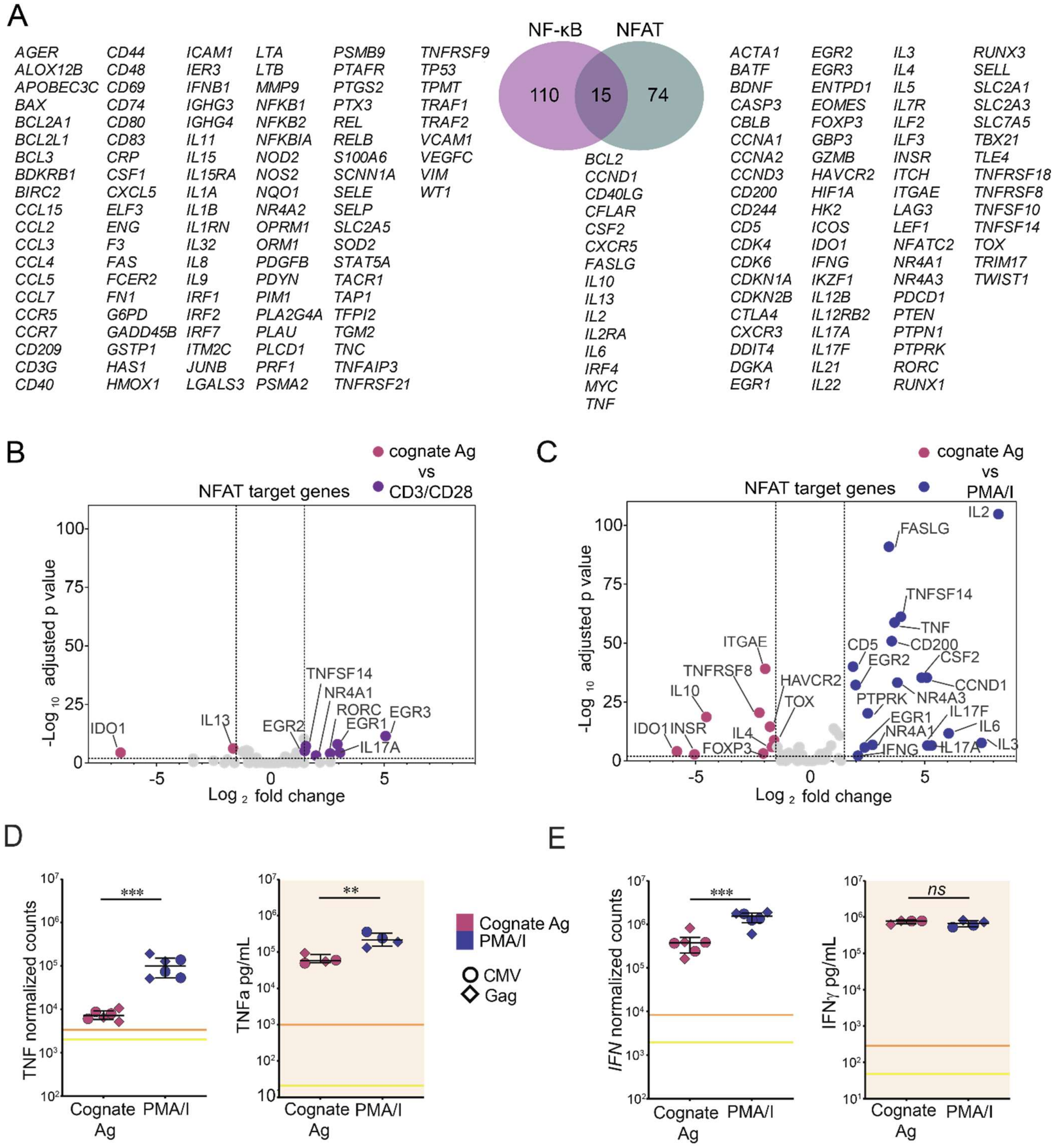


**Supplemental Figure S3**. **Additional analyses on gene and protein expression upon stimulation with cognate Ag versus PMA/I.** (**A**) List of NF-κB and NFAT target genes of interest. (**B**) Volcano plot showing differentially expressed genes (DEGs) target of NFAT in cells stimulated with cognate Ag versus anti-CD3/CD28 (**C**) Volcano plot showing DEGs genes target of NFAT in cells stimulated with cognate Ag versus PMA/I. (**D**) Normalized counts of *TNF* (left) and supernatant levels of TNFα (right) in cells stimulated with cognate Ag versus PMA/I. (**E**) Normalized counts of *IFN* (left) and supernatant levels of IFNγ (right). Supernatant levels measured after 18h stimulation with PMA/I or cognate Ag presented on the surface of autologous DCs. Data presented in units of pg/ml. (**C-D**) Individual symbol corresponds to a single study participant. Horizontal bars represent median with interquartile range All data shown as average of technical replicates. Statistical significance was determined by paired t-test, *P<0.05, **P<0.01, ***P<0.001, ****P<0.0001.

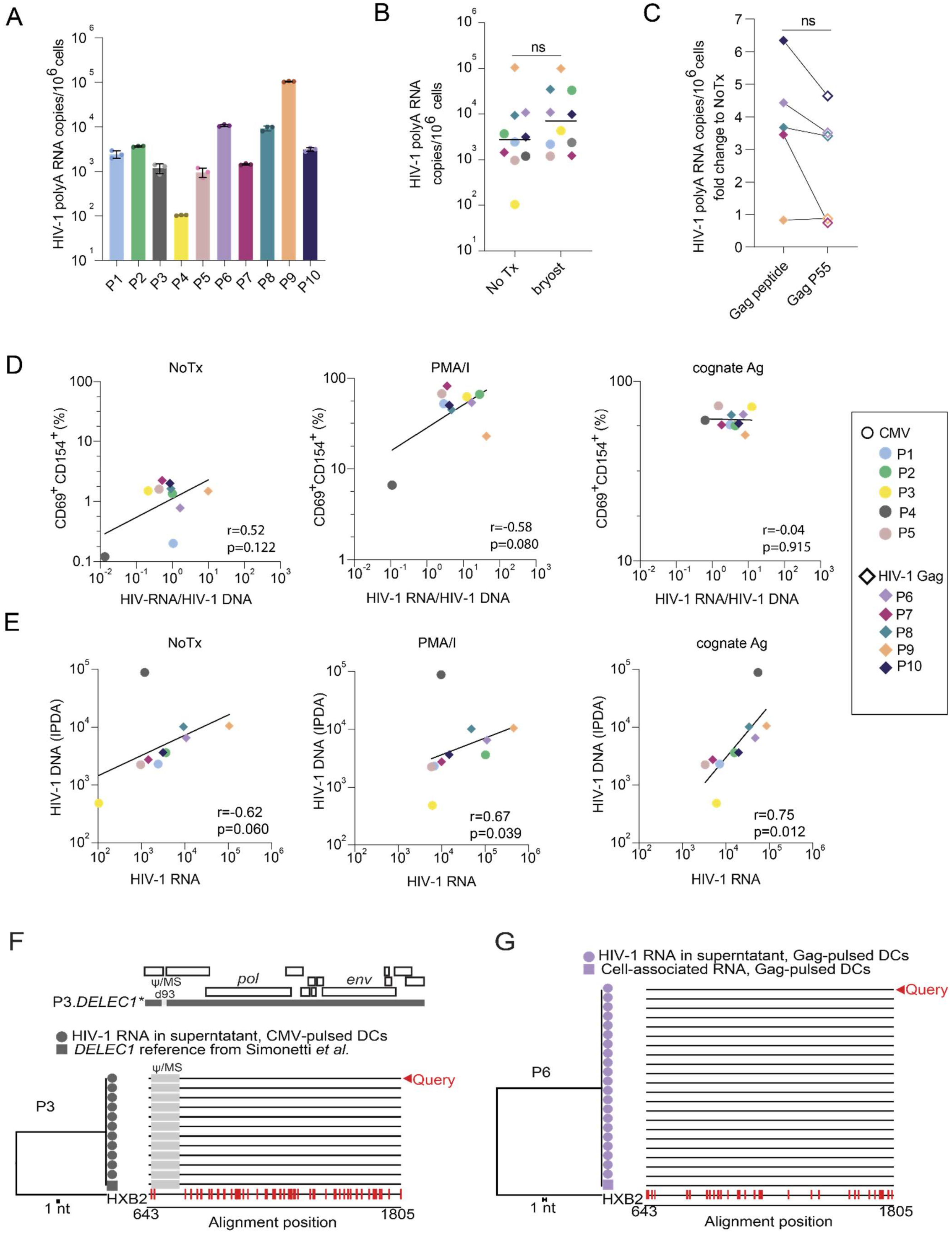


**Supplementary Figure S4. Additional analyses of HIV-1 RNA expression upon re-stimulation. Related to Figure 4.** (**A**) HIV-1 polyA RNA quantification within expanded pool of CMV-responding cells (participants P1-P5) and Gag-responding cells (participants P6-10) for untreated conditions (NoTx) across all study participants. Data shown as triplicates with standard deviation. (**B**) HIV-1 polyA RNA quantification and comparison between NoTx and bryostatin. (**C**) HIV-1 polyA RNA quantification within Gag-responding cells upon stimulation with DCs loaded with Gag peptide pool versus loaded with Gag P55 whole protein. (**B-C**) Individual symbols are shown as mean HIV-1 polyA RNA values for the corresponding study participant and significance was determined by paired t-test. (**D**) Correlation plots of percentage of $CD69^{+}CD154^{+}$ cells and HIV-1 RNA/HIV-1 DNA ratio for NoTx, PMA/I and cognate Ag (CMV/Gag). (**E**) Correlation plots of total number of proviruses as quantified by IPDA and mean HIV-1 polyA RNA for NoTx, PMA/I and antigenic (CMV/Gag) treatment condition group. For **D** and **E**, statistical significance was determined by Spearman's correlation. (**F**) Sequence analysis of virion-associated HIV-1 RNA in supernatant for P3 matched the previously characterized 5'-Leader defective provirus integrated outside of the DELEC1 gene. (**G**) Sequence analysis of virion-associated HIV-1 RNA in supernatant for P6 matched one of the two proviruses contributing to cell-associated RNA. In both **F** and **G**, neighbor-joining trees (based on p-distance, rooted to the HXB2 reference sequence) are flanked by highlighter plots showing mismatches relative to the query.

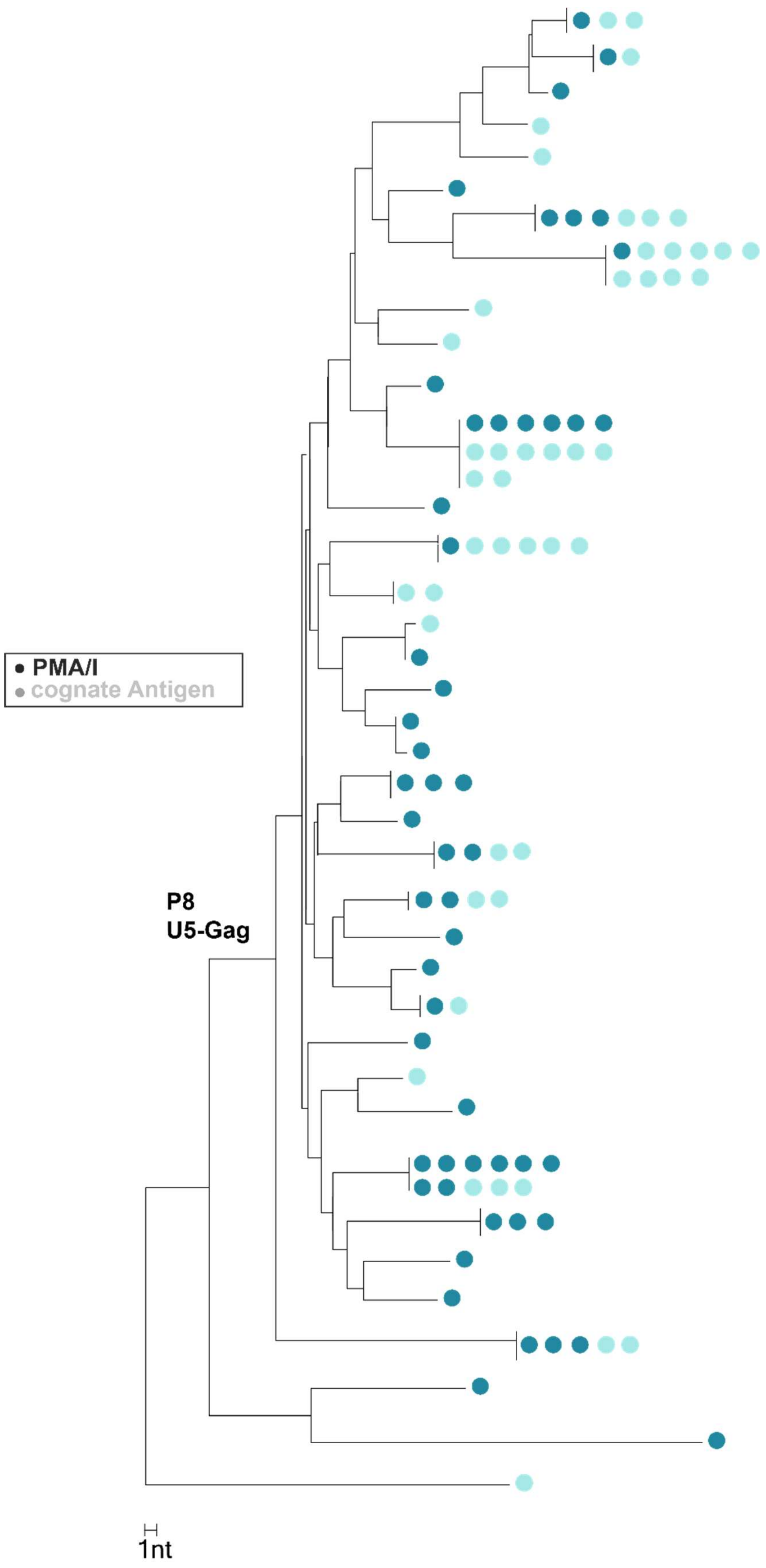


**Supplementary Figure S5**. **Identical HIV-1 RNA sequences are commonly induced both by PMA/I and cognate Ag regardless of the sampling size, related to Figure 5**. Neighbor-Joining tree of HIV-1 U5-gag cDNA sequences from study participant P8.

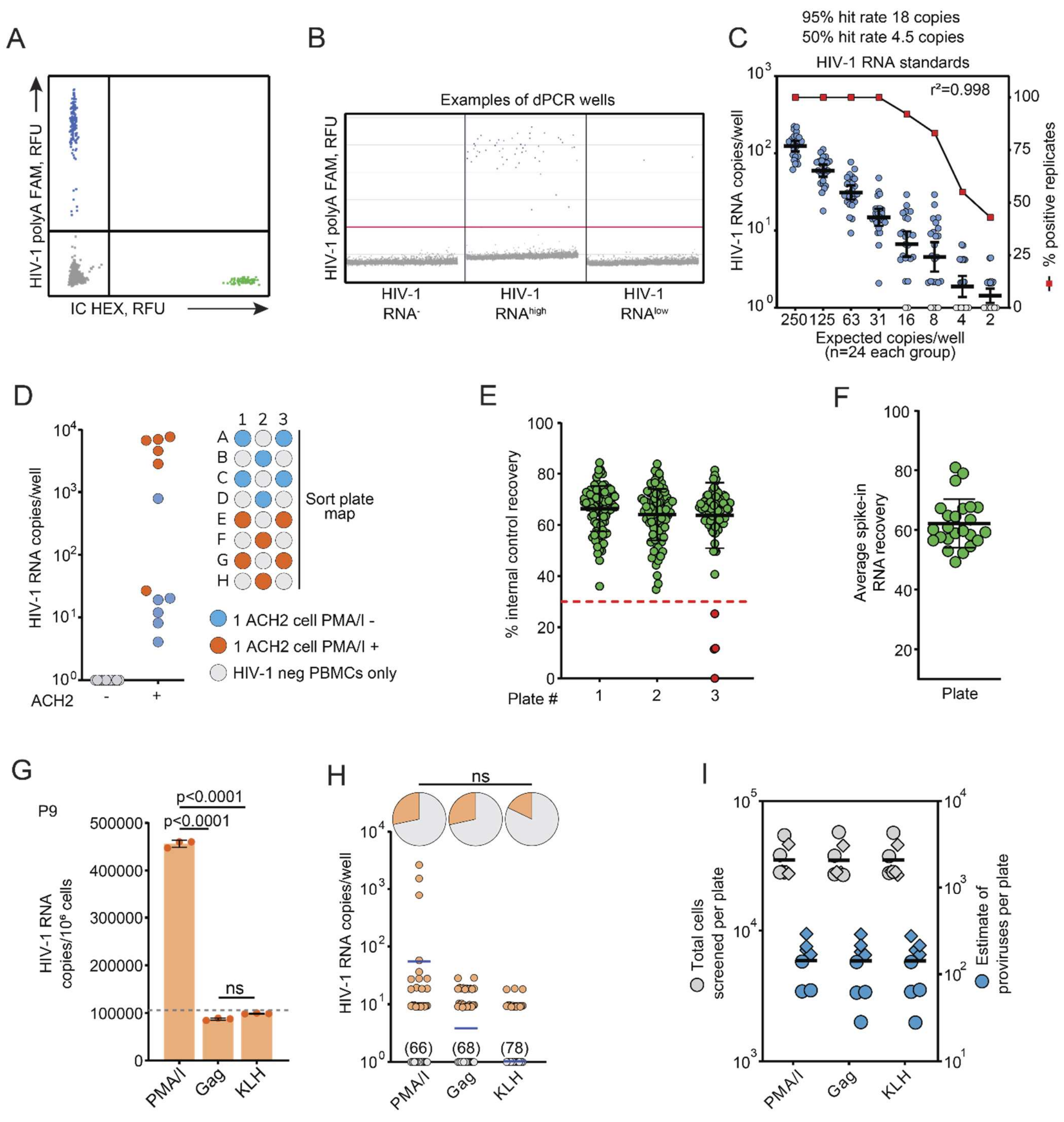


**Supplementary Figure S6. Validation experiments and additional analyses related to the limiting-dilution viral quantification assay. Related to Figure 6.** (**A**) Representative 2-dimentions plot showing positive partitions for HIV-1 polyA RNA in FAM and internal control (IC) in HEX. (**B**) 1-dimention raw data from representative cells with no, high, or low HIV-1 RNA expression. (**C**) Validation experiment to determine the assay sensitivity; known copies of HIV-1 RNA commercial standards were added to wells of a 96 well plates (2-fold dilutions, 24 replicates per dilution); red squares indicate the % of positive replicates. (**D**) Quantification of HIV-1 polyA RNA copies from HIV-1 negative PBMCs or singly-sorted ACH2 cells l with or without PMA/I stimulation. The plate sorting map was designed to assess inter-well contamination and non-specific signal in HIV-1 negative wells. (**E**) Percentage of internal control (IC) recovery from 3 representative plates from the same experiment. Wells with IC RNA recovery lower than 30% IC were excluded from the analysis. (**F**) Percentage of average IC RNA recovery for all 24 participant-derived samples. (**E-F**) Horizontal bars represent mean and standard deviation. (**G**) HIV-1 RNA quantification from bulk cells, as in Figure 4A, for participant P9; the dashed horizontal bars indicate the mean value for cells without treatment (NoTx). (**H**) Limiting-dilution viral quantification assay for participant P9, from the same sample used in **G**; blue horizontal bars indicate mean values; gray symbols indicate values below the limit of detection (9 copies/well) and numbers in parenthesis indicate the number of negative wells; pie charts at the top indicate the percentage of HIV-1 RNA$^+$ wells; the difference between the three groups was assessed by Fisher's exact test. (**I**) Total cells screened in each experiment (grey symbols) and estimate of total proviruses seeded in each per plate (based on total DNA quantification by the IPDA); horizontal bars indicate mean values.

**Supplementary Tables**

| Participant study ID | Age | Sex | Race | Years since HIV-1 diagnosis | Time on ART (years)[a] | Time on suppressive ART (years)[b] | ART Regimen[c] | Ag stimulation |
|---|---|---|---|---|---|---|---|---|
| **P1** | 59 | M | White | 22.8 | 9 | 8 | ABC/3TC/DTG | **CMV** |
| **P2** | 56 | F | AA | 22 | 22 | 15 | ABC/3TC/DTG | |
| **P3** | 57 | M | AA | 24 | 16 | 9.5 | TDF/FTC/RPV | |
| **P4** | 65 | M | White | 28.7 | 28 | 11.8 | TAF/FTC/RPV/DTG | |
| **P5** | 69 | M | Mixed | 27.4 | 27.2 | 16.1 | DES, DRV, RAL, RTV, HGH | |
| **P6** | 60 | M | White | 31 | 26.4 | nsv | TAF/FTC/BIC/FTR | **Gag** |
| **P7** | 63 | F | AA | 16.3 | 14 | 11 | TAF/FTC/ATV/c | |
| **P8** | 42 | F | AA | 28 | 9.4 | 9 | ABC/3TC/DTG | |
| **P9** | 43 | M | AA | 22 | 18 | intermittent nsv | ABC/3TC/DTG | |
| **P10** | 50 | M | AA | 12 | 11 | 11 | TDF,FTC,ATV | |

[a] Years since the beginning of first ART regimen

[b] Years of stable suppression with plasma HIV-RNA below limit of detection of clinical assays

**Supplementary Table S1**. Participant characteristics. Abbreviations: AA (African American), ABC (abacavir); ATV (atazanavir); ATV/c (atazanavir/cobicistat); DRV (darunavir); DTG (dolutegravir); EFV (efavirenz); ETV (etravirine); FTC (emtricitabine); FTR (fostemsavir); nsv (nonsuppressible viremia); RAL (raltegravir); RPV (rilpivirine); TAF (tenofovir alafenamide fumarate) TDF (tenofovir disoproxil fumarate); ZDV (zidovudine); 3TC (lamivudine.

| Primers used for single genome sequencing (SGS) PCR | | | | | |
|---|---|---|---|---|---|
| **Assay** | **Primer name** | **PCR reaction** | **Primer type** | **Primer sequence** | **HXB2 position** |
| *u5-gag* | u5gagSGS_fo | outer | forward | GTARCTAGAGATCCCTCAGAC | 584-604 |
| | u5gagSGS_ro | | reverse | TGACATGCTGTCATCATYTCYTC | 1819-1841 |
| | u5gagSGS_fn | nested | forward | AAATCTCTAGCAGTGGCGCC | 623-642 |
| | u5gagSGS_rn | | reverse | CATCATTTCTTCTARTGTAGCTSCT | 1806-1830 |
| **Primers used for Sanger sequencing** | | | | | |
| **Assay** | **Primer name** | **PCR reaction** | **Primer type** | **Primer sequence** | **HXB2 position** |
| *u5-gag* | u5gagSGS_fn | Sanger | forward | AAATCTCTAGCAGTGGCGCC | 623-642 |
| | INT2A5 | Sanger | forward | GTGATACCCATGTTTTCAGCATTA | 1291-1314 |
| | INT3A2 | Sanger | reverse | AGCTTCCTCATTGATGGTCTCTTT | 1393-1416 |
| | INT3A2_p2 | Sanger alternative | reverse | AGCCTCCTCATTAATGGTATCTTT | 1393-1417 |

| Primers and probes used in bulk HIV-1 RNA ddPCR experiments (5' to 3') | | | |
|---|---|---|---|
| **Forward primer** | **Reverse Primer** | **Probe** | **Fluorophore** |
| CAGATGCTGCATATAAGCAGCTG | TTTTTTTTTTTTTTTTTTTTTTTTGAAGCAC | CCTGTACTGGGTCTCTCTGG | FAM |

| Primers and probes used in limiting dilution HIV-1 RNA assay | | | | |
|---|---|---|---|---|
| **Target Name** | **Forward primer** | **Reverse Primer** | **Probe** | **Fluoro-phore** |
| Internal control (IC) | GAACAGGCCTCGACTTCAAT | TGGTAATGGCAGGTCTAGAAAC | TAAAGCTTGGCTGGTGCATAGGCA | HEX |
| polyA HIV-1 RNA | GCCCTCAGATGCTRCATATAA | TTTTTTTTTTTTTTTTTTTTTTTTGAAG | TGCCTGTACTGGGTCTCTCTGGTTAG | FAM |

**Supplementary Table S2**. Primers and probes used in this study.

| Internal RNA control used for the limiting-dilution HIV-1 RNA assay |
| --- |
| ACACTCTTTCCCTACACGACGCGGGGGTAATACGACTCACTATAGGGGAAATGGACGTTGGTTACCTTCCTGAGTACTGT |
| GGAACATTTCTGCACCCCCATCCGTGGGAACGGCTGAATTGGAAAATGGAACCTACCGAAAGTTCAAAAAAGGGGTTTC |
| TGGACATTTTGCGCTCAGAGCTGGAGTTCACTAAAAAAGAACACTTCACACAATGCATGTCTCACGTTGCGTTATTTTAAT |
| GCATCAAGGAACGTGGGTTAAATCTGCACGGCCGTACATCCAAAAAAATTTAATATCACATGGAAGAAGCAGGCTCCTG |
| AAAAGAAAACGGCAGAAAGGAGAAAAAATCCCGCTGCTGGCATTGGAGCCTGGAACGGAACCCTCGCGCGGTTCAGAC |
| TAAACCCGGATTGTTTAAGACGAACACGGGTACCATCGGTGCGGTATCTCTGGACTTTTCTCCGGGTACAAGTGGATCGC |
| CGATAATCGACAAATGACGTAAGAGTTGTGGATCTTTATGGTGATGGTGTTGCCATAAAGAGTGGAGTATTTGTGAGTG |
| TTATAGCCCCAATTGAAAAGAAGCGCTTAAAACCACTCCCCGTAGCAAAATGACTTTTAGCGAGAGAGAAGATTAACCTT |
| CATGGACTAGCACGCAGGAAAGAGAGCGTCGCAAGGATACATTCCAGCCATGATCCGAGAAACCTTGAAACGGGGTCT |
| GAGAGCACTAAAGCTGGCCGCCGTTAGAGCTGTGGCAGCTGAAGTGGCGTAAACTGAGTGAAGACTTCCAGTGAGATA |
| CAAAACCCCAACCTTCCAAACTTAGCACGTTAGGAGGAAAGTTATGGATCTAATGCGTAATGCCCCGTACTATGCGTCTG |
| CTTTCTCCTGTACGCGTCCCCAATTATAATTTAATCATAATGGATGAGGCAATTTCTCAAACACAGCAGGTTTAACAACTG |
| GAAGACACATCGCAGCCAGAATGGAAATGGGTACGTCAACTGGGGTTCTCATGACAGCCGCTGCCGCGGGAGGTCGAG |
| ATTCACTTCCTTCGTGCGATGCACCAGAGTTAAATGAAAAAGGAAAAATCGCTGAACGTCCATGGCATTCTGGATATGAG |
| ATCTCGCATCTAACGGGAAGATTGTCTTGTTCGTTCTAAGCATAAAAAGCAGGAAATGATTTGACAACTGGCGTAACAGT |
| TGATGGAACGTAAGTAATACAACTCAGTAGGACGCCTGCTGATCCTTTCGATGAGTCGTCTTGAGCCGATGATCGGTATC |
| AGCTGGCTTCAGTTGACCTTCCAGAAATGGGTGCCTATTAGTAAACTGCGTGGACTGTGAACACCGGACGCTGCCTAAAA |
| CCAATCATATTAACAGATGGTTCGCTCAGGGTAATTTTGGCAGGATCTGTGCCAGTAACACAGTAGTGCCGCGCAGCGTA |
| GAGGGCGGATTGGCCGGAATCCGAACCAAATGAAGATGACATCAACCTATACTTGGGAAAACTTATGGAAGATGATGA |
| AAACAGTGCATACCGGTAAAAAGCCTAAGTGCTCGTAAATTACGTTGACGCACCAGAAAGAGTCATTTTTAGTATGCCTG |
| AACCAAAGTGTTAAAAAATGGATGCCACTTATGGCGAACACGGTGTAAGAAGAAAAGCAAGGAAAACCTCTGTGGACCT |
| AATAAGGCGAAGAAACTTATCAAAGTGGATGGTTAACCAAATGGCAGCTTAAAGCTAGCACTATGCAGACTGAAGGCG |
| GTGTGTTAATGGAACTGAGAACTTGTTAAGACCTGCTGTAACCAACAGGTGCGAAAAGAGTGCAGCGCATTTGCAGCCG |
| TAAGAAAGTCAGTAACCCGGAGCCTAATCACCGTAACGGCTAAGCAAGCAATATACATGACTCTGAAGACAAACGTCGC |
| ACGGGGCAGCCCAGTTCAGCAGCCCATGGCTGAAGCCGTTGGAAGGGCGTGCAGAGTTGCTCACAGTGAACAGCAGGA |
| GATCCACGGGATACTGCATTTACAGACACCTCAGGCTACAGCCATCGTCGCGATTTGAGCTTATTCTTGATGTCAGGGCG |
| GGGTATAGGTAAGATGACTTTGGGTATGTGCTGTATTATCACGGCATCCATTCTGTTGTGGTACGCACAAATACAGCCTC |
| ACTGGATTGCCGCAAGTATTATTTTGGAATTCTCTCTGGTAAGCAGAGAACAGCACCCCGGCCATCGTCCGACTTCACGC |
| ATAAGTTTGGATGCAGTTTGTAGAGGCTACAATATCCGTTACACAAACGCTGAGACGTGACACTAAGAATTCCTGTCTGA |
| ATGTATACCGAATTCGTATTCGCAGCCATCTAATTCTGTGAATGGGTCCCGCGAGCGTATGGCGATAGTAAAGGATGGA |
| CACCGCTCTAGCCCATCCCGTCACTGGGGTAAGGGTGGCATCGTGACATGTGCCATGTTCAAATGTAAAAAGAATATGG |
| AAGGAAAGATTGTACAACCTGAGAATCTGGAATATACGATTGTAGTAACGCCACATAGTGGTGAGGAGCACGCGGTGG |
| GAAATGATACAGGCAAACACGGCAAAGAAATCAAAGTTACTCCTCAGTCATCCATTACTGAAGCTGAATTAACCGGTTAC |
| GGAACAGTAACAATGGAATGCTCACCCAGAACAGGCCTCGACTTCAATGAGATGGTGTTGCTGCAGATGGAAAATAAAG |
| CTTGGCTGGTGCATAGGCAATGGTTTCTAGACCTGCCATTACCATGGCTCTTCAAA |

**Supplementary Table S3**. Sequence of the internal RNA control used for the limiting dilution HIV-1 RNA assay.
